# Resilience of A Learned Motor Behavior After Chronic Disruption of Inhibitory Circuits

**DOI:** 10.1101/2023.05.17.541057

**Authors:** Zsofia Torok, Laura Luebbert, Jordan Feldman, Alison Duffy, Alexander A. Nevue, Shelyn Wongso, Claudio V. Mello, Adrienne Fairhall, Lior Pachter, Walter G. Gonzalez, Carlos Lois

## Abstract

Maintaining motor behaviors throughout life is crucial for an individual’s survival and reproductive success. The neuronal mechanisms that preserve behavior are poorly understood. To address this question, we focused on the zebra finch, a bird that produces a highly stereotypical song after learning it as a juvenile. Using cell-specific viral vectors, we chronically silenced inhibitory neurons in the pre-motor song nucleus called the high vocal center (HVC), which caused drastic song degradation. However, after producing severely degraded vocalizations for around 2 months, the song rapidly improved, and animals could sing songs that highly resembled the original. In adult birds, single-cell RNA sequencing of HVC revealed that silencing interneurons elevated markers for microglia and increased expression of the Major Histocompatibility Complex I (MHC I), mirroring changes observed in juveniles during song learning. Interestingly, adults could restore their songs despite lesioning the lateral magnocellular nucleus of the anterior neostriatum (LMAN), a brain nucleus crucial for juvenile song learning. This suggests that while molecular mechanisms may overlap, adults utilize different neuronal mechanisms for song recovery. Chronic and acute electrophysiological recordings within HVC and its downstream target, the robust nucleus of the archistriatum (RA), revealed that neuronal activity in the circuit permanently altered with higher spontaneous firing in RA and lower in HVC compared to control even after the song had fully recovered. Together, our findings show that a complex learned behavior can recover despite extended periods of perturbed behavior and permanently altered neuronal dynamics. These results show that loss of inhibitory tone can be compensated for by recovery mechanisms partly local to the perturbed nucleus and do not require circuits necessary for learning.

## Introduction

All animal species display behaviors crucial for reproduction and survival that are challenged by a changing environment. These behaviors rely on activity in brain circuits with an appropriate balance between excitatory and inhibitory neuronal activity (E/I balance). Once disturbed, brain circuits strive to restore E/I balance^1^as its loss leads to abnormal neuronal firing patterns, which are implicated in several brain diseases, including epilepsy^1,2^. In this study, we explore how brain circuits react to extensive and long-term perturbation of inhibition.

Zebra finches produce a highly stereotyped song with minimal variability over extended periods^3^, underpinned by temporally precise neural activity^4^. To chronically perturb the E/I balance, we genetically blocked inhibitory neurons in HVC, a premotor brain nucleus of the male zebra finch involved in song production. HVC contains two main types of excitatory neurons that project into two downstream targets: Area X (proper name) and nucleus RA. HVC also contains inhibitory neurons whose axons do not leave HVC^4^. Previous work has shown that acute pharmacological blocking of interneurons in the HVC of adult finches perturbs song for a few hours until the chemical is washed out, after which normal behavior is restored^5^.

Transient perturbations do not help us understand the outcome of prolonged E/I imbalances, such as long-term adaptations that the system might undergo. To investigate whether an adult motor circuit can overcome extended periods of perturbation, we used viral vectors carrying the light chain of tetanus toxin to chronically silence inhibitory neurons in the HVC of adult zebra finches. The persistent loss of inhibitory tone altered brain activity and severely perturbed song production for several weeks, during which animals produced degraded, variable songs. Most surprisingly, the song completely recovered around two months after being perturbed. While the song was degraded, we observed increased expression in MHC I and microglia marker genes within HVC, suggesting the potential involvement of inflammatory and immune signaling pathways in neural circuit reorganization.

To study how the chronic silencing of interneurons affects circuit dynamics, we performed acute and chronic recordings in HVC and its downstream target, nucleus RA. Surprisingly, although the neuronal activity within HVC or RA remained abnormal even two months after the perturbations, the song was fully restored. In addition, in adult animals subjected to perturbation of interneurons in HVC, the song degraded and recovered after lesioning LMAN, a song nucleus essential for song learning in juveniles^6–8^. This indicates that the mechanisms by which song recovers in adults after perturbation differ from those used by juvenile animals to learn the song.

Our findings indicate that an adult motor circuit can regain the original complex learned behavior without restoring the original neuronal dynamics after extended periods of drastic E/I imbalance. Together with our previous work perturbing HVC excitatory neurons, these observations suggest that loss of either inhibitory or excitatory tone in a brain nucleus can recover by mechanisms that are likely in part local and homeostatic since they do not require practice or inputs from nuclei necessary for song learning such as LMAN.

## Results

### Long-term song degradation after genetic muting of local inhibitory neurons in the zebra finch

To functionally mute interneurons, we used the light chain of tetanus toxin (TeNT), which blocks the release of neurotransmitters from presynaptic terminals, thereby preventing neurons from communicating with their postsynaptic partners^9^. We used an AAV viral vector expressing TeNT under the human dlx5 promoter, which is selective for HVC inhibitory interneurons^5,10^. The virus was injected stereotaxically into the HVC of adult male zebra finches. The expression of TeNT does not directly alter the ability of interneurons to fire action potentials but effectively prevents synaptic release, thereby muting them. As a control, a second group of animals was injected with an AAV carrying the green fluorescent protein NeonGreen driven by the ubiquitous cytomegalovirus (CMV) enhancer fused to the chicken beta-actin promoter (CAG). Throughout various time points after virus injection, we recorded songs, measured changes in gene expression at single-cell resolution, and obtained electrophysiological measurements (chronic and acute within HVC).

An average male zebra finch song contains 4 to 7 units called syllables. These syllables are fixed in order and make up a so-called motif. The motif is repeated multiple times to form a bout ^9–11^, defined as the set of vocalizations produced by a bird without pauses (> 200 msec) between them. Throughout all experiments, we continuously monitored the songs of individually housed animals from 7-20 days before virus injection to 60-100 days post-injection (dpi) using sound-isolated chambers. In control animals, the song did not change at any time after the viral injection (Figure 2d). For the first 24-48 hours, the song of TeNT-treated animals was identical to their original song. However, starting at 2-5 dpi, the song of TeNT-treated animals began to degrade, both in its acoustic and temporal features (Figure 1a). By 5-10 dpi, the song of TeNT-treated animals was highly degraded and bore no resemblance to the original song before viral injection (Figure 1a).

**Figure 1:**
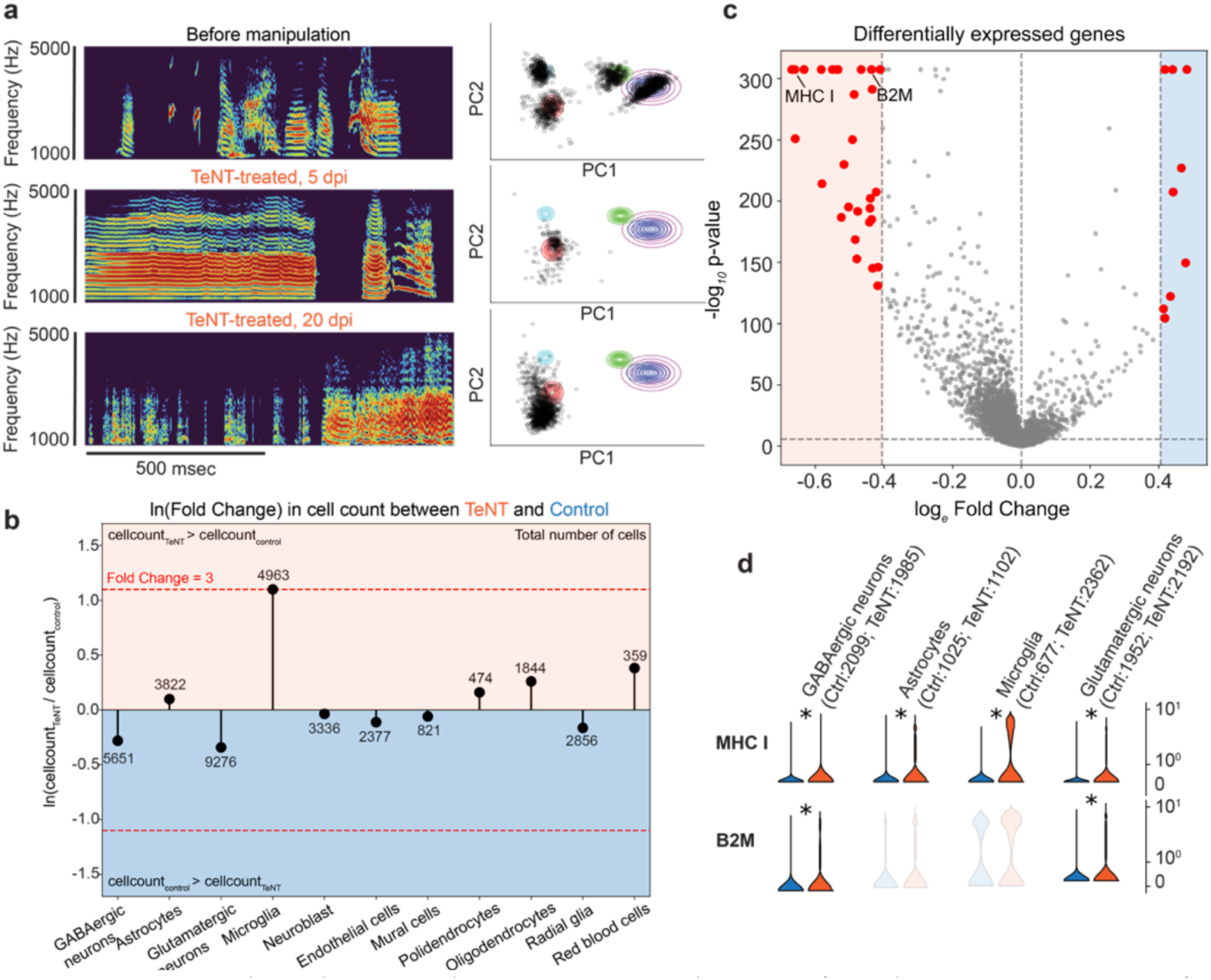
Song degradation and transcriptomic changes after chronic inactivation of inhibitory neurons in HVC by viral expression of tetanus toxin (TeNT). **a (left)** Examples of spectrograms of songs produced by an animal treated with TeNT before, and 5, 20 days post-injection (dpi). **a (right)** 2D principal component analysis (PCA) visualization of syllables aka song segments produced by the animal on the same day as the spectrogram example on the left. Note: even when the songs are degraded, we continue to refer to an uninterrupted length of vocalization as a ‘syllable or song segment’. Individual dots indicate single syllable renditions. In the unperturbed song (before virus injection), syllables cluster into distinct groups (each differently colored circle represents a distinct syllable in the PCA space). **b** Log-fold change in total number of cells per cell type between TeNT-treated and control animals. **c** Volcano plot showing statistical significance over magnitude of change of differentially expressed genes between TeNT-treated and control animals across all cell types. Dotted lines indicate fold change = 1.5 and p value = Bonferroni corrected alpha of 0.05. **d** Violin plots of normalized counts of major histocompatibility complex 1 α chain-like (MHC1) (ENSTGUG00000017273.2) and beta 2 microglobulin-like (B2M) (ENSTGUG00000004607.2) genes in control (n=2, blue) and TeNT-treated (n=2, red) animals per example cell cluster. A star indicates a significant increase in gene expression in TeNT-treated animals compared to control (p < 0.05 and fold change > 1.5).

**Figure 2:**
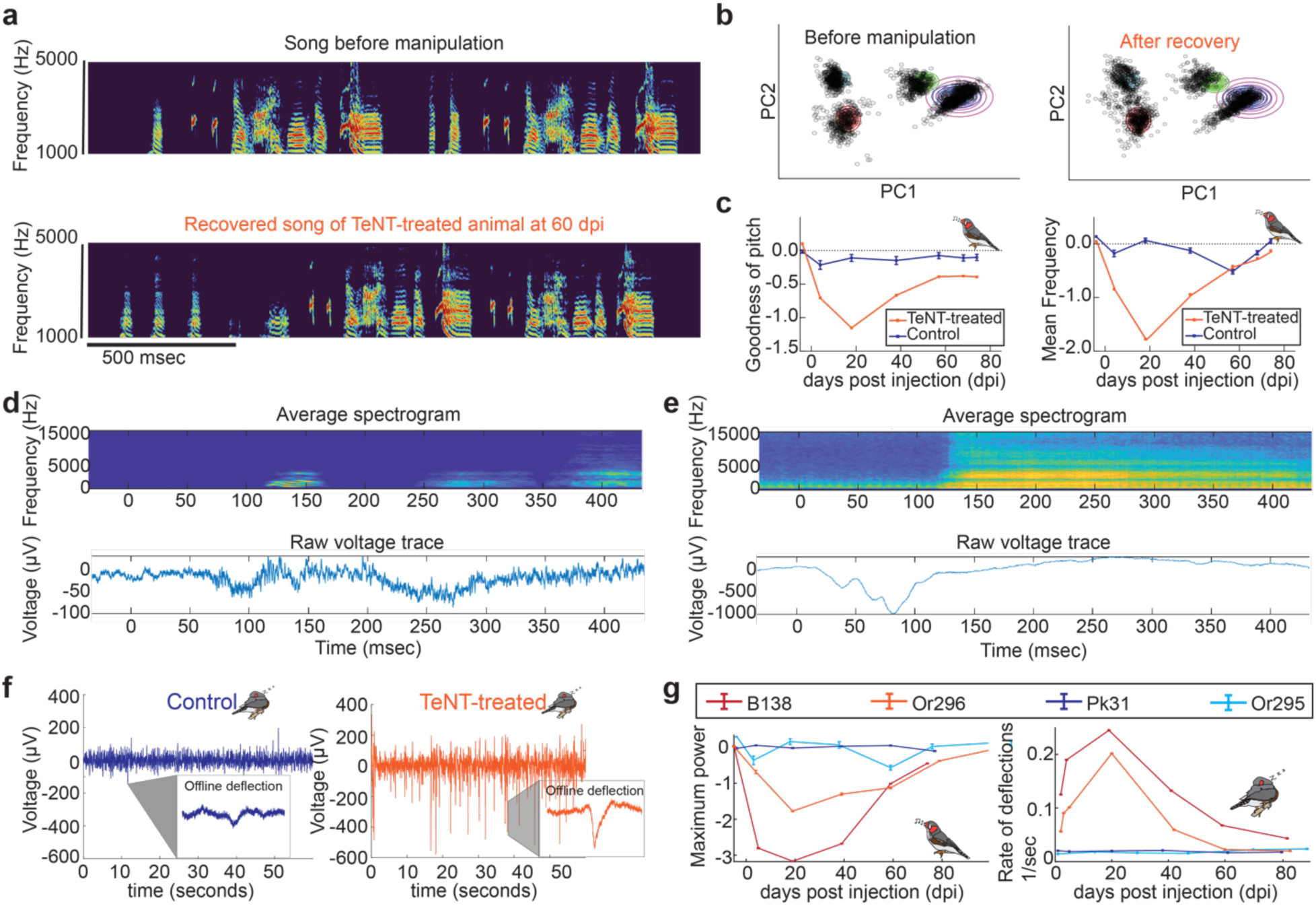
Behavioral and chronic electrophysiology recordings in HVC after chronic inactivation of inhibitory neurons by viral expression of TeNT. **a** Examples of spectrograms of songs produced by an animal treated with TeNT before, and 60 days post-injection (dpi). Data from this animal (V648) are also shown in c. **b** 2D principal component analysis (PCA) visualization of syllables aka song segments produced by the animal on the same day as the spectrogram example in **a**. **c** Different features of the vocalizations averaged/grouped before and after stereotaxic injection of control (GFP-expressing) or TeNT virus. The grouped mean frequency and goodness of pitch measures represent averages over song segments. Control animals (Or295 and PK31) are labeled in blue (average of n=2); TeNT-treated animals (V648, B183, B138 and Or144) are labeled in orange (average of n=4). Traces indicate averages over syllables sung within the 3 closest days of recorded vocalizations. Error bars indicate standard error. Values are normalized relative to distributions 5 days before perturbation (see Methods). **d (top)** Average spectrogram of 5 vocalizations in a control animal **d (bottom)** Average raw deflection traces aligned to the vocalizations. **e (top))** Average spectrogram of 5 vocalizations of a TeNT-treated animal **d (bottom)** Average raw deflection traces aligned to the vocalizations**. f** Raw electrophysiology traces during night time (lights off period) from TeNT-treated and control animals 5 days post-injection (dpi). Note: electrodes were implanted on the same day of viral injection. Insets show examples of stereotypical deflections that at 5 dpi were approx. 50 ms in duration, and - 530 µV in amplitude, in TeNT-treated animals, compared to 96 ms and -140 µV in control animals. **g (left)** Quantification of song quality of chronically implanted animals over time. Control animals (PK31 and Or195) are depicted in shades of blue. TeNT-treated animals (B138 and Or296) are depicted in shades of orange and red. **g (right)** Rate of deflection events during night time. Control animals (PK31 and Or195) are depicted in shades of blue. TeNT-treated animals (B138 and Or296) are depicted in shades of orange and red. The highest rate of sleep voltage deflections in TeNT-treated animals occurred during the periods when the songs were most degraded, as shown on the left.

During the first few days of song degradation, TeNT-treated animals produced instances of syllables with abnormally long durations far outside the range of any adult syllable duration observed in control animals (Figure 1a shows one such vocalization at 5 dpi; examples of syllable length distribution are shown in Supplementary Figure 1). By 8 to 30 dpi, vocalizations became very short (< 100 msec) with inter-syllable intervals of varying lengths (100 msec to 1 sec) (spectrogram examples in Supplementary Figure 2). At 20-30 dpi, the song of TeNT-treated animals was highly variable between renditions and lacked the typical organization of motif and bout (Figure 1a).

In previous work, we showed that silencing of projecting neurons in HVC using a similar approach induced a transient song degradation with full recovery after 2 weeks^11^. Based on this finding, we anticipated that the time course for song recovery should be similar when perturbing the inhibitory neurons of the same circuit. However, even 4 weeks after genetic muting of HVC interneurons, the song did not recover. Although inactivation of projection neurons leads to song degradation and recovery in two weeks, the loss of inhibitory tone in HVC did not result in song recovery within a month of perturbation, suggesting permanent behavioral damage.

### Chronic interneuron inactivation in adult songbirds increases the expression of MHC I and the number of microglia in HVC

To understand the molecular mechanisms caused by interneuron muting, we performed single-cell RNA sequencing (scRNA-Seq) of HVC from control (n=2) and TeNT-treated (n=2) adult male zebra finches at 25 dpi, around the time of peak song distortion. HVC from both hemispheres of all four birds were dissected based on retrograde tracer fluorescence (injected retrograde tracer into area X) and dissociated to prepare single-cell suspensions, which were indexed and pooled. This allowed the construction of a combined dataset containing results from all animals and conditions without the need for batch correction (Supplementary Figure 10, Supplementary Table 2). Following quality control, we retained a total of 35,804 single-cell profiles from four animals.

While cell type abundance was highly concordant between replicates of the same condition (Supplementary Figures 10, 16A), we found that TeNT-treated animals displayed a three-fold increase in the number of microglia (Figure 1b). In addition, we found a significant increase in the expression of the α chain of MHC I and β2-microglobulin (B2M) genes across several neuronal cell types in TeNT-treated animals (Figures 1c and d). This increase in microglia and MHC I was not due to an inflammatory reaction caused by the surgical procedure or the AAV injection since control animals also received a viral injection with an identical serotype viral backbone that did not affect neuronal activity. Thus, we hypothesize that the increase in microglia and MHC I in TeNT-treated animals results from the chronic muting of inhibitory neurons.We also confirmed the increase in MHC I and microglia by *in situ* hybridization (ISH) (Supplementary Figures 12, 13).

Previous works have shown that MHC I and microglia are involved in the synaptic reorganization of brain circuits, especially during postnatal development in mammals^12–18^. In addition, several studies have demonstrated the upregulation of microglia following seizure-like activity in mammals^19,20^. Our finding suggests that the increase of microglia and MHC I expression caused by chronic silencing of interneurons may be involved in the synaptic reorganization of HVC.

### Near-perfect behavioral recovery after prolonged zebra finch song degradation

Silencing HVC interneurons drastically degraded the song, leaving its structure unrecognizable within 2-3 weeks. Notably, the song remained highly variable even three weeks after the manipulation, with differences observed between renditions produced on the same day. (Supplementary Figures 1). This is unlike the abnormal vocalizations observed after other irreversible song circuit manipulations, such as tracheosyringeal nerve resection or deafening, which result in song production that eventually becomes degraded but stable between renditions ^21–23^.

We wondered whether the songs from animals in which we permanently inactivated inhibitory neurons would eventually stabilize into a fixed degraded song or whether the song would keep changing without ever stabilizing. To this end, we decided to record their behavior for an extra 4 weeks. Unexpectedly, after 40-70 days of degraded song production, the original song started to re-emerge. By around 70 dpi, the song of all TeNT-treated animals was highly similar to the song produced before viral injection (Figure 2a).

Next, we asked whether the transcriptomic changes observed at 25 dpi persisted after song restoration. To this end, we performed ISH using a probe against RGS10, a microglia marker gene, at 25 dpi (when the song is fully degraded) and 90 dpi (after the song has recovered). At 25 dpi, the number of RGS10+ cells significantly increased in TeNT-treated animals compared to control animals (Supplementary Figures 12 A, 13 A, and B) and returned to normal levels by 90 dpi.

To ask whether long-term loss of inhibitory tone results in similar transcriptomic changes in HVC as during song learning, we counted the number of RGS10+cells in HVC at different times during song learning in naive (untreated) juvenile birds using ISH. The number of microglia in the HVC of juveniles was high during the early stages of song learning (20-50 days post-hatching (dph)) and decreased after that (Supplementary Figures 12 B, 13 C, and D). These observations suggest that microglia might be involved in fine-tuning brain circuits to achieve appropriate function during song learning and behavioral recovery following perturbation. However, the increase in microglia after silencing inhibitory neurons in adult animals could account for two opposite scenarios. The increased inhibitory tone in HVC and the number of microglia could induce synaptic changes that contribute to degraded song production. Alternatively, the rise in microglia could be part of the recovery response to produce synaptic changes needed to regain the song following perturbation.

### Chronic muting of interneurons in HVC results in long-term abnormal neuronal dynamics that recover in parallel with song quality

Next, we sought to understand the neuronal dynamics within HVC during behavioral perturbation and recovery due to the loss of inhibitory tone. Towards this end, we continuously recorded the behavior and neuronal activity in HVC for 90 days in freely moving animals after injection of interneuron muting (TeNT) or control viral vectors.

We implanted a 4-shank electrode array containing a total of 16 recording sites in both in TeNT-treated and control animals. We simultaneously collected electrophysiological data, head movement (recorded with the accelerometer built into the head stage), and audio recordings for 90 dpi. We found that single units were detectable for only 24-48 hours, after which the signal irreversibly degraded. However, these electrodes allowed us to record local field potential (LFPs) for several weeks. LFP recordings capture local field oscillations caused by synaptic events and action potentials from many neurons and neurites in a volume around the electrode^24^. We analyzed the spectral decomposition of the LFPs during song production from degraded vocalizations in TeNT-treated or control animals (Figures 2d and e, Supplementary Figures 6 A and B). As expected from prior work^25,26^, both the song behavior and LFP features remained stable in control animals: the electrical activity during singing showed voltage deflections of small amplitude, which were present throughout the song (Figure 2d, Supplementary Figure 6 B). In contrast, at 5 dpi, TeNT-treated animals produced abnormally long vocalizations that were preceded by large voltage deflections, not present in control animals (Supplementary Figure 6 A). Between 10 and 70 dpi, the songs of TeNT-treated animals were highly variable, both between renditions and across days (Supplementary Figure 1). Due to this behavioral instability, the vocalizations of perturbed animals could not be reliably aligned across trials. To find a stable measure allowing to compare neuronal activity between control and experimental groups, we focused on electrophysiological events occurring during nighttime (lights-off periods when the animals are not moving or singing) (Figure 2f, 2g (right)). The neuronal activity recorded during the nighttime period will be referred to as “offline” or “lights-off” throughout this manuscript.

HVC activity during sleep is hypothesized to be a form of replay of neuronal firing similar to that occurring while the animal sings^27–30^. Therefore, analyzing electrophysiological activity in HVC during offline periods throughout the perturbation could reveal disruptions related to degraded behavior. As observed during singing, during offline periods, the voltage deflections seen in TeNT-treated animals were more frequent and of larger amplitude than in the control group (Figure 2f, Supplementary Table 1B). At 5 dpi, the average amplitude of the voltage deflections was around 3-fold higher in TeNT-treated than in control animals (-530 µV and -140 µV in TeNT-treated and control animals, respectively), with a 280% increase in TeNT-treated animals (Supplementary Table 1). In addition, in TeNT-treated animals, these large amplitude deflections at night occurred more frequently when the song was most degraded (Figure 2g right panel). In TeNT-treated animals, the rate of deflections returned to those seen in control animals as the song regained stereotypy around 50 dpi (Figure 2g).

Offline deflection events were further characterized by spectral decomposition of the LFPs in the 1-200 Hz frequency range. At 5 dpi, we observed increased power only in the low gamma band (30-70 Hz) in TeNT-treated animals, which returned to control levels as the songs regained stereotypy (Supplementary Figure 6 C and D). These findings suggest a relationship between behavioral recovery and the return to control-like offline electrophysiological activity, supporting the hypothesis that offline activity may correlate with the quality of song execution.

### Offline neuronal activity in HVC remains abnormal even after the song recovers

To examine local neuronal activity at single-neuron resolution during song degradation and recovery, we performed acute head-fixed recordings. High-density silicon probes (Neuropixels)^31,32^ allowed the simultaneous recording of neurons in HVC, RA, and intervening brain areas (Figure 3a). As described above, we obtained the most reliable measurements of abnormal brain activity in TeNT-treated animals during offline periods, given the variable song production. Therefore, we performed Neuropixel recordings in head-fixed animals when the lights were off during their natural sleep cycle. We focused on acute recordings obtained within a few hours after probe insertion to achieve the best possible signal-to-noise ratio. To understand how neuronal activity evolves as the song degraded and recovered, we performed acute recordings at 3-6, 20-22, and 70-77 dpi in TeNT-treated and 10 and 30 dpi in control animals. To control for potential artifacts caused by the surgery or viral injection, we included recordings from one naïve animal (not injected with any virus).

**Figure 3:**
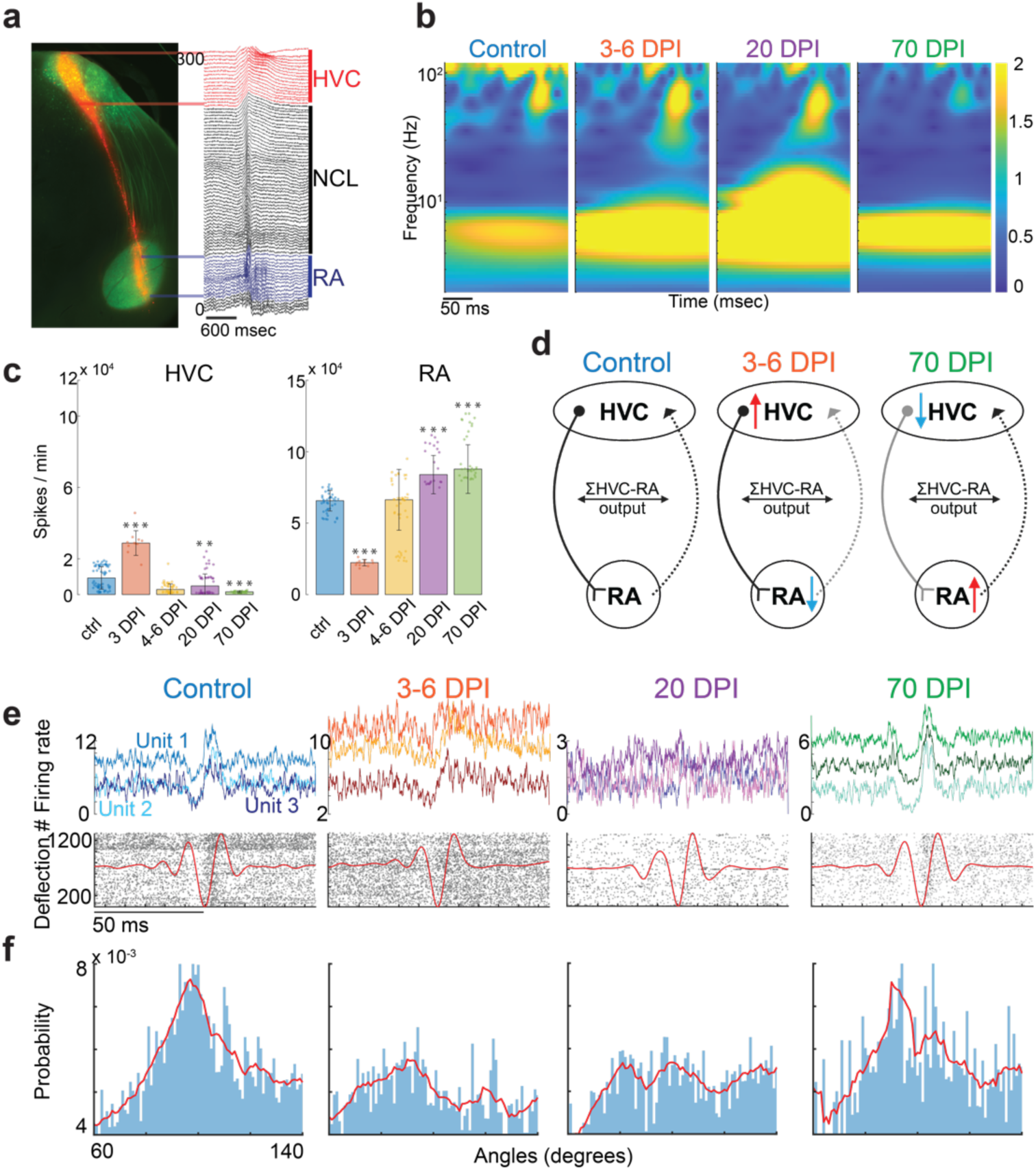
Acute electrophysiology recordings after chronic inactivation of inhibitory neurons in HVC by viral expression of TeNT. **a** Histological section example of a brain slice from a control animal shows the track of the probe (NPIX coated with DiI) insertion (red) and GFP from the viral injection (green). The raw electrophysiology traces are color-coded by brain area (HVC in red, NCL in black, RA in blue). **b** Spectral decomposition of LFPs of the average normalized deflections during lights-off acute recordings in control animals and TeNT-treated animals at 3-6, 20, and 70 dpi. **c** Average firing rates of neurons within HVC and RA in control animals (n=3) and TeNT-treated animals at 3 (n=1), 4-6 (n=4), 20 (n=4) and 70 (n=4) dpi. In HVC, the increase in average firing rate at 3 dpi after TeNT injection significantly differs from control (p-value from rank sum test: 0.4*10^-6^), as well as the decrease in average firing rate after TeNT injection at 20 dpi (p = 0.002) and 70 dpi (p = 0.57*10^-11^). In RA, the decrease in average firing rate at 3 dpi significantly differs from control (p = 0.43*10^-6^), as well as the increase in average firing rate at 20 dpi (p = 0.09*10^-9^) and 70 dpi (p = 1.45*10^-13^). The stars above the bar plots indicate statistical significance (* = p < 0.005, ** = p< 0.01, *** = p < 0.001). **d** Model of HVC-RA circuit dynamics before and after TeNT-perturbation in HVC based on c. **e (top)** Example traces of the firing rate of three individual neurons during deflection events that lock or do not lock to a specific phase of the alpha oscillation (shown in red in bottom plots) in HVC of control animals and TeNT-treated animals at 3-6, 20, and 70 dpi. **e (bottom)** Examples of neuronal firing (indicated by black dots) for one neuron per condition in HVC. **f** Probability distribution of the phase of firing events relative to the alpha oscillation of the LFP in control animals and TeNT-treated animals at 3-6 dpi (n=4), 20 dpi (n=4), and 70 dpi (n=2).

Loss of inhibitory tone led to seven events resembling seizure-like activity in HVC (n=3 TeNT-treated animals at 3-20 dpi), which propagated downstream into the nucleus RA (Supplementary Figure 9 B). These seizure-like events in HVC resemble the bursts of increased activity previously reported in rodents after acute pharmacological blockade of interneurons or during seizure events^1,2,33^ We investigated spontaneous offline activity to understand the overall HVC and RA dynamics after perturbation. For the first 3-4 dpi, HVC started with a heightened level of spontaneous offline activity (likely due to the loss of inhibition). However, after 20 dpi, activity decreased to a level below that of control animals (Figure 3c). Surprisingly, the activity in HVC in TeNT-treated animals remained substantially reduced relative to the control animals (*p = 0.57*10^-11^*) (Figure 3c) even after song recovery. While activity in HVC in TeNT-treated animals was initially elevated and eventually decreased, RA showed the opposite trend (Figure 3c). This observation is consistent with a prior study^34^ showing acute changes in RA firing patterns in single neurons upon stimulation of HVC in brain slices. These observations suggest that HVC may directly increase the activity of RA interneurons. According to this model, the initial increase in overall activity in HVC (caused by blocking of HVC interneuron activity) would activate interneurons in RA, thereby reducing overall activity in RA (Figures 3c and 3d). In contrast, when the overall activity in HVC was permanently reduced, the activity in RA increased to keep the overall sum of HVC-RA output constant (Figures 3c and 3d). However, it is essential to note that our recordings in RA do not allow us to identify whether the recorded activity originated from excitatory or inhibitory neurons.

### Degradation of behavioral stereotypy is paralleled by loss of local neuronal firing precision to specific phases of alpha oscillations

We used acute Neuropixel recordings to understand the relationship between neuronal activity and LFP oscillations during offline voltage deflection events, which seem to change on the same time scale as the behavioral recovery. This readout could explain the relationship between input into a circuit, the local neuronal dynamics^24^, and the execution of the recovered behavior.

We found acute offline voltage deflections similar to those we observed with our chronic recordings (Figure 2f and Supplementary Figure 7). We detected comparable changes during the perturbation (Figure 3b) as in the chronic recordings (Supplementary Figure 6 C and D). We observed significant differences in the LFP signatures of voltage deflections between TeNT-treated and control animals, namely in the alpha (1-10 Hz) and the low gamma ranges (30-40 Hz) (Figure 3b). Previous studies have suggested that local neuronal dynamics cause low gamma oscillations and that synaptic inputs may cause alpha oscillations^24^. This could mean low gamma oscillations represent local neuronal activity, while alpha is a readout of inputs into HVC, highlighting their importance in circuit dynamics.

Next, further confirming the findings of the chronic recordings, acute Neuropixels recordings revealed that after an initial increase in the power of the low gamma range (30-40 Hz), it returned to control levels around 70 dpi (Supplementary Figure 9 A). The phase relationship between alpha and gamma oscillations was variable during song degradation but returned to resemble controls after song recovery (Supplementary Figure 9 C-E). In contrast to the gamma oscillations’ power, alpha oscillations stayed significantly higher in TeNT-treated animals, even after the song recovered at 70 dpi (Supplementary Figure 9 A).

To further study the relationship between local neuronal activity and LFP dynamics, we compared the timing of single neuron firing with the alpha and low gamma LFP phase during offline deflections. We employed methods previously used to characterize sharp-wave ripple events during offline activity in the hippocampus^35^. The neuronal firing within HVC locked tightly to specific phases of the alpha and low gamma LFPs in control animals (Figures 3e and f; Supplementary Figure 8A). This phase-locking relationship remained unchanged in RA and in the local neuronal precision locked to a specific phase of gamma LFPs within HVC during the interneuron manipulation (Supplementary Figure 8). However, the precision of local neuronal firing to alpha oscillations between 60 to 140 degrees was lost or broadened during the song degradation period (Figure 3e and f). The animals’ songs started to recover at the same time as the local neuronal firing in HVC locked again to a specific phase of the alpha oscillations (Figure 3 e and f).

In conclusion, we found that restoration in behavioral stereotypy was accompanied by the return to normal of several aspects of neuronal dynamics, such as the low gamma LFP power and spike-locked alpha oscillation in HVC. However, even after the song recovered, several features of offline HVC dynamics remained abnormal, including alpha LFP power in HVC and spontaneous activity both in HVC and RA. These observations indicate that the adult brain can overcome extended periods of abnormal neuronal activity and perturbed behavior and precisely restore a complex behavior without recovering the original neuronal dynamics. This suggests that HVC may transition into a new state capable of producing the original behavior despite the persistence of altered neuronal dynamics.

### Song degradation and recovery in adults do not require a brain nucleus essential for juvenile learning

Our transcriptomic results suggested that some mechanisms might be shared between song learning in young animals and recovery after inhibitory circuit manipulation in adults. To test this hypothesis, we investigated whether LMAN, a brain nucleus essential for song learning, is involved in song degradation and recovery after interneuron manipulation in adults.

Juvenile animals start singing a highly variable and unstructured song. By continuous trial and error over 2-3 months and thousands of practice renditions, their song “crystallizes” into a stable motif with minimal variability^36,37^. Song learning in juveniles requires LMAN, another nucleus in the song system that provides input to RA, the downstream target of HVC^6–8^. During learning, LMAN injects noise into RA to enable the exploration of different states^38^ and gates the plasticity of synapses from HVC to RA^39^. In addition to its role in learning, LMAN is necessary for song modification in adult animals^40^. Deafening adult animals causes a progressive degradation of their songs over several weeks, which can be blocked by prior LMAN lesioning. A previous study reported that adult HVC microlesions led to days-long song degradation that was preventable by prior lesioning of LMAN^41^. Therefore, it is possible that the ‘noisy’ input from LMAN to RA becomes more dominant when HVC is perturbed in adult animals. This noisy LMAN input into RA could underlie the production of degraded songs after manipulation of HVC activity.

To test whether the change in LMAN input is the main cause of degraded song production (given our findings about decreased HVC activity) and is necessary for song recovery after interneuron perturbation, we chemically lesioned LMAN by bilateral ibotenic acid injection, and 10 days later, we injected AAV-dlx-TeNT into HVC (n=3). The song degraded and recovered similarly in all TeNT-treated animals, regardless of the presence of an intact or lesioned LMAN. This observation indicates that song degradation after HVC interneuron perturbation is not simply due to an exacerbated LMAN influence on RA (Figure 4 & Supplementary Figure 3-5). In addition, previous works have demonstrated that the modification of the dynamics of RA requires input from LMAN, both in juveniles and adults^42,43^. Our results revealed that perturbation of HVC caused profound changes in the activity of RA during song degradation and recovery. Thus, the ability of the song to recover after LMAN lesioning suggests that in adult animals, RA can modify its dynamics without the participation of LMAN. These findings indicate that nuclei of the song system play different roles during adult song degradation and recovery following HVC perturbation and juvenile song learning^7,44^.

**Figure 4:**
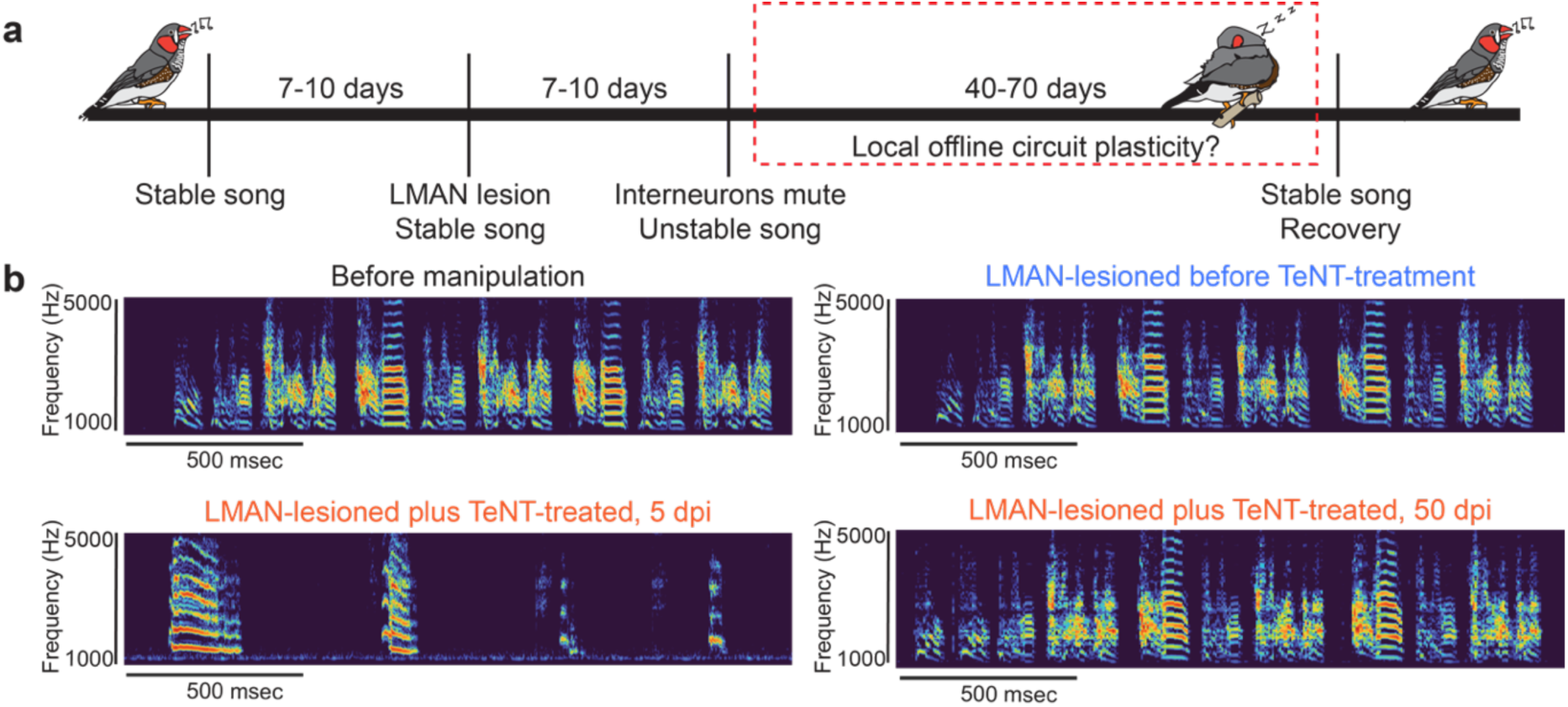
Song degradation and recovery after interneuron perturbation without LMAN a brain area necessary for song learning. **a** Schematic of the experimental design. We injected adult male animals n=3 with ibotenic acid to lesion LMAN 7-12 days before the TeNT-virus injection into HVC and recorded the song before lesion, after lesion and after interneuronal manipulation. **b** Spectrogram example of the song of an adult male zebra finch showing the behavioral readout of the schematic presented in **a.** N=3 adult males degraded and recovered their song after interneuron perturbation in HVC without LMAN (LMAN lesion was confirmed by CGRP immunohistochemistry shown in Supplementary Figure 3-5 along with spectrogram examples)

## Discussion

The brain must balance excitation and inhibition to maintain the appropriate levels of physiological activity needed to execute reliable behaviors. We found that permanently muting inhibitory neurons in a pre-motor circuit in adult songbirds severely disrupt their song for an extended period (40-70 days). However, despite this long-term disruption, animals eventually recovered and produced a version of the song that was highly similar to the original.

To understand the cellular mechanisms underlying this behavioral recovery, we investigated the transcriptomic and electrophysiological changes induced by this perturbation. Our sc-RNA sequencing data revealed a threefold increase in microglial marker genes compared to control animals, which returned to normal levels as the song recovered. In mammals, microglia are known to play a role in synaptic changes during the assembly of brain circuits in the embryonic and perinatal brain^17,18,45^. Consistent with their suggested role in developmental plasticity, we observed an increase in microglia during song learning in juveniles. Additionally, seizures have been shown to elevate the number of microglia^19,20,46^. Our findings suggest that microglia could participate in reorganizing HVC after chronic perturbation of inhibitory neurons. However, whether microglia aid or delay the recovery process remains an open question for future studies.

To understand how prolonged perturbation of interneuron activity affected brain dynamics, we performed long-term electrophysiological recordings in HVC during both song degradation and recovery. We detected changes in the electrophysiological properties of offline events, which are known to be important for a replay of the song in HVC. During song degradation, we observed an increase in the rate and amplitude of these deflection events, which returned to normal levels once the song regained its stereotypy. To investigate neuronal dynamics with greater resolution, we analyzed single and multi-neuronal activity within HVC and its downstream nucleus RA in acute recordings with Neuropixel probes. Following the muting of interneurons in HVC, we observed two persistent changes in HVC dynamics that remained abnormal even after the song had fully recovered: (i) a decrease in offline single neuron firing and (ii) an increase in alpha frequency power. Alpha frequency oscillations have been suggested to result from synaptic inputs into a brain region^24^. Therefore, we posit that the decreased neuronal firing in HVC may have initiated a compensatory increase in the synaptic inputs originating from upstream nuclei projecting into HVC (such as Uva and Nif). This could account for the observed increase in alpha power. Alternatively, this increase in alpha power could originate from synaptic inputs from neurons within HVC. However, we believe this scenario is unlikely because we found decreased local neuronal activity in HVC. Future experiments will be necessary to understand how the perturbations of HVC affect the dynamics of other nuclei in the song system and the cell types within HVC.

To investigate the relationship between LFP oscillations and local neuronal dynamics in HVC, we measured the precision of single neuronal firing to specific phases of alpha and gamma oscillations, called spike locking. We observed significant changes upon the muting of inhibitory neurons. In control animals, HVC neuronal firing locked to alpha and gamma oscillations during offline voltage deflections. In contrast, single neuronal firing did not lock to alpha frequency oscillations during the peak song distortion. As the song recovered, the spike locking to alpha (input-driven) oscillations returned to normal.

The relationship between single neuronal firing and alpha oscillations in the song system is not yet understood. We hypothesize that this relationship could represent how the local HVC circuit responds to inputs from other brain areas. In this scenario, during song degradation, when stereotypy of the song is lost, the local HVC circuit cannot correctly respond to synaptic inputs from upstream nuclei (mainly Nif and Uva) in the song system, resulting in the observed desynchronization of spikes to alpha oscillations. As the song recovers, spike-field coherence becomes more precise even though the power of the alpha oscillations remains permanently elevated. These findings suggest that the motor circuit can execute behavior properly despite permanent changes in its neuronal dynamics.

We observed superficial similarities between juvenile song learning and adult song recovery after HVC interneuron perturbation, such as variable song production and increased microglia numbers. However, a fundamental difference exists between these processes. In juveniles, song learning requires input from LMAN to RA, and it has been hypothesized that LMAN injects noise into RA, allowing HVC inputs into RA to explore different states^6,7^. Previous studies have also shown that LMAN is required to enable song modifications in adults^43,47^. Here, we found that in adult animals without LMAN, the song degraded following the inactivation of inhibitory neurons in HVC and recovered after several weeks. Further studies are necessary to find both the commonalities and differences in mechanisms of brain reorganization following HVC perturbation in adults and the development of the song circuit in juveniles.

In our previous work, we observed that the song was drastically degraded and recovered after permanent genetic muting of a large fraction of projection neurons in HVC in adult animals^11^. In the present study, we observed that chronically inactivating a large fraction of interneurons in HVC also causes song degradation and eventual recovery. Our results reveal that HVC can produce reliable songs, a complex learned behavior, after significantly reducing the fraction of its active excitatory or inhibitory neurons. These studies highlight this nucleus’s unique resilience by demonstrating that HVC can restore behavioral dynamics after two “orthogonal” perturbations to the circuit. Our current and previous studies show that extensive perturbations of HVC can recover by mechanisms that are likely, in part, local and homeostatic since they do not require practice or inputs from nuclei necessary for song learning, such as LMAN. Furthermore, our findings indicate that behavior can be recovered after a severe and prolonged brain E/I balance perturbation without restoring the initial dynamics. Instead, it appears that HVC adapts and reorganizes into a new state capable of producing the original behavior. This adaptive process may be crucial for maintaining motor behaviors over long periods and could represent a key feature of brain resilience.

## Methods

### Animals

The Institutional Animal Care and Use Committee of the California Institute of Technology approved all procedures involving zebra finches. All birds used in the current study were bred in our own colony and housed with multiple conspecific cage mates of mixed genders and ages until used for experiments. Before any experiments, adult male birds (>120 days post-hatching (dph)) were singly housed in sound isolation cages with a 14/10 hr light/dark cycle for >5 days until they habituated to the new environment and started singing. Thereafter, birds were kept in isolation until the end of the experiment.

### Behavioral recordings

Adult male zebra finches’ (n=30, 130-890 dph) undirected songs were recorded 24/7 in sound-isolated chambers for 10-14 days before any manipulation to get a baseline of their song. Recordings were done with microphones (Audio-technica, AT831b) that are connected to an amplifier M-TRACK 8 and recording software Sound Analysis Pro 2011 at 44100 Hz. Animals were housed in these chambers and continuously recorded for the duration of the experiments.

### Viral vectors

AAV-TeNT contained the promoter from the human dlx5 gene driving expression of the light chain of tetanus toxin fused to EGFP with a PEST domain. AAV9-dlx-TeNT was obtained from the Duke viral core facility. The control virus used was AAV9-CAG-NeonGreen, where CAG drives the expression of NeonGreen, which is a GFP variant.

### Stereotaxic injection

Birds were anesthetized with isoflurane (0.5% for initial induction, 0.2% for maintenance) and head-fixed on a stereotaxic apparatus. First, to inject a retrograde tracer in area X, craniotomies were made bilaterally and fluorescent tracers (fluoro-ruby 10%, 100-300 nL) were injected through a glass capillary (tip size ∼25 μm) into the corresponding nuclei (coordinates from dorsal sinus in mm – area X: Anteroposterior (AP) 3.3-4.2, Mediolateral (ML) 1.5-1.6, Deep (D): 3.5-3.8). To deliver the virus (AAV) into HVC, a second surgery was performed 7-10 days after retrograde tracer injection. By then, HVC was strongly labeled by fluorescence and visible through a fluorescent stereoscope. AAVs diffuse extensively (∼500 µm), and a single injection (∼100 nL) in the center of HVC was sufficient to label enough cells. All injections in HVC were performed at ∼20 nL/min to minimize physical damage. At the end of every surgery, craniotomies were covered with Kwik-Sil, and the skin incision was closed with Gluture.

#### LMAN lesion

We used ibotenic acid (2%) dissolved in 100mM NaOH to chemically ablate LMAN, in adult male zebra finches (n=4) between 120-200 dph. Coordinates used with 50 degree head angle were (3.2/3.5/3.8 mm AP; 1.7/1.8/2 mm ML; 2.4;2.8/2.4;2.7/2.3;2.5 mm Deep). We achieved ablation of ∼80-98% of LMAN bilaterally confirmed by CGRP staining. We initially measured the acoustic variability within each syllable before and after lesion and found that it decreased in 3/4 animals (which agrees with previous findings) to confirm the success of our LMAN lesions. Next, 7-10 days following the LMAN ablation we followed the previously describe stereotaxic injection protocol for tracer and virus injection. The one animal that did not have decrease in syllable level acoustic variability following lesion later on by histology was also confirmed to only receive a partial lesion therefore we removed this animal from further behavioral analysis.

### Chronic electrophysiology recordings

Animals (n=4, 300-700 dph) were implanted in the right hemisphere HVC with 4 by 4 electrode arrays (Neuronexus A4x4-3mm-50/100-125-703-CM16LP) based on retrograde fluorescent labeling of HVC (just as for viral injections). Post-perfusion histology images were obtained to locate the electrode array within HVC for each animal (Supplementary Figure 15). Electrode implantation occurred within the same surgery as the viral injection. This procedure follows the same surgical steps as the viral delivery protocol, until the point of electrode implantation. A small opening was cut on the dura (just big enough to fit the electrode array) to lower the electrodes manually. The reference and ground were a gold screw pin placed into the cerebellum. The skin was removed from the surface of the skull for the majority of the surface, in order to secure the implant. Before implantation, the skull and the craniotomies were cleaned with saline and dried and the skull was prepared according to the protocol of the C&B Metabond cement system. Post implantation we covered the craniotomies with kwik-sil. Once hardened, we covered the whole skull, and the part of the electrode still exposed, with metabond. The head stage (Intan RHD Part # C3335) was connected to the probe before implantation and securely metabonded to the connection between the probe and head stage in order to prevent detachment when the bird is moving. SPI interface cables (Intan Part #C3203, #C3213) were connected to the acquisition board (Open Ephys). Data was recorded at 30,000 Hz with the Open Ephys software system. Animals were freely moving with a passive counterweight-based commitator system.

### Acute electrophysical recordings

Animals (n=10, 140-250 dph) went through the same surgical procedure as described for a stereotaxic viral injection. However, at the end of the surgery the skin was removed from the skull, and the whole skull was pre-treated and covered in metabond except for the craniotomies over HVC that were covered with kwik-cast until the day of the acute recording session. Shortly before the recording session, a head-bar was glued on top of the frontal surface of the metabonded skull to allow the head-fixation of the bird for the recording session. Then, the kwik-cast was removed from the craniotomy over HVC (left or right hemisphere or both depending on the animal) and a small incision was made in the dure over HVC, which was identified by the retrograde tracer previously injected. The ground was placed into the cerebellum. Then the high-density silicone probe (Neuropixel) was lowered with a motorized arm over hours for 2.6-3 mm deep into the brain. The head stage and acquisition board was connected to the computer and data was recorded with the Open Ephys software. Once the probe settled in the brain, we had 4 distinct recording sessions. Post-perfusion histology images were obtained to locate electrode array within HVC for each animal (Supplementary Figure 16). Recording sessions: lights on silence (10 min), followed by playback of the bird’s own song (3-10 min); lights-off silence (10 min), followed by playback of the bird’s own song (3-10 min); microinjection of 100 nL 250 µM Gabazine (Hellobio, HB0901), followed by the same protocol of lights-off and on without Gabazine.

### Song analysis

Song analysis was performed using MATLAB (MathWorks).

#### Song feature parameterization

Continuous audio recordings (44.1 kHz) were segmented into individual bouts manually. We used the open-source Matlab software package, Sound Analysis Pro 201^48^ to generate spectrograms and derive non-linear, time-varying song parameterizations. The time-varying features were: pitch, goodness of pitch, Wiener entropy, log-power of the spectrogram, mean frequency, and the maximum power in each quadrant of the frequency range 0-11 kHz (labeled power 1, power 2, power 3, and power 4). These features were computed across the entire bout every 1 ms. These time-varying features were the base from which various portions of the song were further characterized.

#### Syllable parameterization

To quantify the change in the acoustic structure of the song, we tracked song features over the course of the perturbation. In Fig 2, we took the average feature values over all song segments in the three closest days of recorded song for each day plotted. We extracted the average mean frequency, average pitch, maximum goodness of pitch, maximum log power, minimum entropy, and entropy variance over the first 50 ms of each song segment, or the entire song segment if the segment was less than 50 ms. We centered and normalized feature values by the mean and standard deviation of the distribution of feature values in syllables five days pre-perturbation to compare trends across birds.

To visualize how the population of song segments evolved relative to the original syllable structure (Fig 1A and 2), we plotted song segments as points in the first 2 principal components of the 7-dimensional acoustic feature space composed of: average mean frequency, average pitch, maximum goodness of pitch, maximum log power, minimum entropy, and entropy variance over the first 50 ms of each song segment, and the full duration of each song segment. 500 syllables recorded pre-perturbation were labeled by hand, and contour plots were made at the mean of each labeled stereotyped syllable type for visual reference as the song degrades and recovers. Principal component dimensions were computed on the 500 labeled syllables, and subsequent song segments were projected into this principal component space.

#### Syllable segmentation

We identified syllables and silences within each bout by imposing thresholds on the time-varying, total log-power of the spectrogram. We selected the threshold based on manual inspection for each bird. We then performed a smoothing step wherein periods of silence less than 15 ms were converted to song segments. Song segments less than 20 ms were excluded from the analysis. Segmentation was further checked by eye by random sampling across both stereotyped motifs and degraded song. We then applied the power threshold to the entire course of song recordings for each bird.

A note on terminology: we refer to song segments to indicate continuous periods of singing. In the unperturbed song, these song segments are termed syllables. Because this is a continuous recovery process, these terms sometimes overlap in our usage.

#### Syllable timing distributions

Syllable durations were extracted from the syllable segmentation process. Discrete distributions of syllable durations were computed by normalizing each distribution of all song segment durations within individual days such that the sum of each daily distribution over all binned durations equaled one. Distributions for individual days were then assembled into a matrix wherein the columns represented normalized distributions for individual days. This matrix was plotted as a heat map in Supplementary Figure 1.

#### Continuous representation of bout trajectory

We generated continuous visualizations of bouts across the entire perturbation trajectory as shown in the graphical abstract^49,50^. We randomly sampled 100 bouts from single days of recording to build a representative sample of the song over the course of the experiment. For each bout, we slid a 150 ms window in 3 ms steps along the bout length. We then generated a high-dimensional, acoustic parameterization of each 150 ms song window by taking the moving average in 20 ms segments every 5 ms of six song features (mean frequency, pitch, goodness of pitch, power 4, power 2, and summed log power of the spectrogram). We performed principal component analysis on this high-dimensional set across all bouts to reduce the feature set to 30 dimensions.

### Analysis of the chronic electrophysiology recordings

#### Preprocessing of Data

Bad channels were removed from the analysis by visual inspection of the first minute of each recording. Three of the four birds studied had one channel that had behavior that was an order of magnitudes larger than all others, and data from this channel was removed for further analysis. Furthermore, channels that appeared out or close to the edge of HVC based on histology were also eliminated from further analysis.

#### Detection of sharp waves

Accelerometer and electrophysiology data were resampled to 125 Hz from the original 30,000 Hz using the Matlab resample function. Since accelerometer data was not zero-centered, the mean was subtracted before resampling and then added back to avoid edge effects from the lowpass filtering during resampling. Sharp waves were detected from the downsampled data using a procedure adapted from a zebra finch sleep study^51^ and were only detected during periods of low movement. This was determined by setting a threshold on the moving standard deviation of the velocity as movement events appeared as large fluctuations in the velocity data. Briefly, the local field potential (LFP) traces were averaged across all electrodes and bandpassed from 1 to 40 Hz using MATLAB’s bandpass function. Putative events were detected by peaks in the inverted (-1x) trace and were in the lowest 5% of all values during non-movement, were at least 10 ms long, and were 80 ms apart. These putative events were used to construct a template by taking the average 80 ms trace centered on each event. Cross-correlating this template with the entire overnight LFP trace and taking the z-score (mean and SD from only non-movement times) revealed events that closely matched the template. Peaks were detected in the z-scored cross-correlation and defined as events if they were above a given threshold and were at least 80 ms apart. Notably, thresholds were not tuned across birds or recordings but a constant threshold was sufficient in all birds to detect events. Extraction of sharp waves. The detected events were used to index the original 30,000 Hz data. 500 ms of data was extracted for each event and recentered on the minimal value in the center 100 ms. An equal number of 500 ms non-sharp wave events were extracted from a random selection of times without movement or sharp waves. Sharp waves with clear electrical noise were omitted from further analysis.

#### Multi-unit activity (MUA)

MUA was detected using an established method^52^. Briefly, channels were common averaged referenced, bandpassed from 600 to 6,000 Hz, and upsampled to 50,000 Hz. The square root of the local energy operator^53^ was taken and then Gaussian smoothed (σ= 0.4). Thresholding this trace and then downsampling back to 30,000 Hz gave the times of high MUA.

#### Time-frequency analysis

The continuous wavelet transform was used to compute the spectra for each sharp wave. Percent increase relative to mean power spectra were averaged within each day and bird. The spectrum of average power in the center 33 ms of each event was used to compute the percent increase in power relative to non-sharp wave events at each frequency. To determine the presence of phase-amplitude coupling between low-frequency (10-20 Hz) and gamma-frequency (30-40Hz) bursts, the phase of the low-frequency band, and power of the high-frequency band were extracted using the Hilbert transform of the band-passed signal. Determining whether the peak gamma activity occurred at a consistent phase of the low-frequency oscillation, could suggest phase-amplitude coupling. To further assess this, we asked at what time and frequency a max in the spectrum occurred for each event, relative to the minima of each sharp wave. We also wanted to ask whether the power of deflections in different frequency bands differed significantly across control and experimental birds. We computed the relative increase in power in the frequency bands of interest (15-30 Hz, and 30-70 Hz) during a deflection relative to non-deflection times. We then used a t-test to statistically test whether power differed significantly across birds for a given day.

### Analysis of the acute electrophysiology recordings

The low-frequency electrophysiology data was recorded at 2,500 Hz while the high-frequency data was recorded at 30,000 Hz. A MATLAB script was written to analyze the low- and high-frequency oscillations. All low-frequency data was high pass filtered (1-300 Hz) and median subtracted. All high-frequency data was common average referenced and low-pass filtered (300-7,000 Hz). For deflection event detection, we used two methods. One to use a template per animal for extraction of deflection and non-deflection events with template matching in a semi-automated manner. Second, manually picked out peaks that were 9 standard deviations away from the average raw signal, then further manually curated these events. Most of our further analysis was done using the deflection events that resulted from template matching (except Supplementary Figure 10A for the difference in power between deflection and non-deflection event calculation). For spike detection, we used two separate methods, one to threshold the high-frequency data to only consider spikes that are 9 standard deviations above the average signal in a 2-minute window. The second method used for spike detection was using the Kilosort software v2.5^54^ to extract spikes from the raw data. Next, we manually curated the dataset to capture the waveform and firing rate of neurons. The thresholded spiking data was used to generate spiking related figures. We performed statistics (Wilcoxon, rank sum test) to show statistically significant firing rates related to control in HVC and RA in Figure 3B.

To calculate the power difference between deflection and non-deflection events in the acute recording in Supplementary Figure 11 A, we used a script developed for the chronic recordings with the use of the built-in MATLAB function *pwelch*. To analyze the spectral decomposition of the LFP signature (Figure 3D) in control and TeNT-treated animals during the experimental timeline we calculated the Morse continuous wavelet transform using the built-in MATLAB function. From this finding, we obtained the mean oscillation at the alpha range (1-10 Hz) and at low gamma (30-70 Hz). Then, we performed an iterative alignment process of these two oscillations and calculated the angle of the low-frequency oscillation (from obtaining the Hilbert transform) at the peak amplitude of the high-frequency oscillation (Supplementary Figure 11BC). Then we proceeded to align the single neuronal firing (obtained from Kilosort) to the alpha and the low gamma oscillations and plotted the normalized probability distribution (Figure 3F) of neurons firing at a specific phase (angle) of the alpha or low gamma oscillations. We performed the Kolgomorov-Smirnov test on the cumulative density function (CDF) (Supplementary Figure 11D) to assess if the change in probability distribution in the relationship between alpha and low gamma oscillations is statistically significant from control distributions at 3-6, 20, and 70 dpi.

### Single-cell RNA sequencing

#### Animals

All of the work described in this study was approved by California Institute of Technology and Oregon Health & Science University’s Institutional Animal Care and Use Committee and is in accordance with NIH guidelines. Zebra finches (Taeniopygia guttata) were obtained from our own breeding colony or purchased from local breeders.

#### Dissociation and cDNA generation

Animals were anesthetized with a mix of ketamine-xylazine (0.02 mL / 1 gram) and quickly decapitated, then the brain was placed into a carbogenated (95% O2, 5% CO2) NMDG-ACSF petri dish on ice. The brains were dissected on a petri dish with NMDG-ACSF surrounded by ice under an epifluorescent microscope guided by the fluoro-ruby retrograde tracing from Area X to HVC.

We used the commercially available Worthington Papain Dissociation system with some minor changes and add-on steps. We followed all the steps included in the Worthington protocol with a final concentration of 50 U/mL of papain. To match the intrinsic osmolarity of neurons in zebra finches we used NMDG-ACSF (∼310 mOsm) instead of the EBSS for post-dissection and STOP solution.

Another modification was to add 20 µL of 1 mg/mL Actinomycin D (personal communication from Allan-Hermann Pool^55^) into 1 mL of the post-dissection medium and the STOP solution in which trituration occurred. Papain digestion occurred for an hour on a rocking \surface with constant carbogenation in a secondary container above the sample vial at RT. We performed trituration with increasingly smaller diameter glass pasteur pipettes. Trituation was performed inside the papain solution. Then, once the tissue was fully dissociated, we centrifuged the samples at 300 g RT for 5 minutes and resuspended them in STOP solution. Next, we used a 40 µm Falcon cell strainer pre-wet with the STOP solution and centrifuged again at 300 g RT for 5 min. Finally, we resuspended the cell pellet in 60µl of STOP solution and proceeded to barcoding and cDNA synthesis. The cell barcoding, cDNA synthesis, and library generation protocol were performed according to the Chromium v3.1 next GEM single cell 3’ reagent kits by Jeff Park in the Caltech sequencing facility. Sequencing was performed on an Illumina Novaseq S4 sequencer with 2x150 bp reads.

#### Generation of count matrices

The reference genome GCA_003957565.2 (Black17, no W) was retrieved from Ensembl on March 20, 2021 (http://ftp.ensembl.org/pub/release-104/gtf/taeniopygia_guttata/). We quantified the gene expression in each of the four datasets using the kallisto-bustools workflow^56^. The reference index was built using the kb-python (v0.26.3) *ref* command and the above-mentioned reference genome. Subsequently, the WRE sequence was manually added to the cdna and t2g files generated by kallisto-bustools to allow the identification of transgenic cells. The count matrix was generated for each dataset using the kallisto-bustools *count* function. The resulting count matrices were compared to those generated by the 10X Cell Ranger pipeline (v6.0.1) and kallisto-bustools *count* with *multimapping* function. For all four datasets, kallisto-bustools mapped approximately 10% more reads than Cell Ranger (Supplementary Figure 16). No increase in confidently mapped reads was observed when using the multimapping function, indicating that reads align confidently to one gene in the reference genome (Supplementary Figure 16).

#### Quality control and filtering

The datasets were filtered separately based on the expected number of cells and their corresponding minimum number of UMI counts (Supplementary Figure 10). Following quality control based on apoptosis markers and library saturation plots (Supplementary Figure 10), the count matrices were concatenated and normalized using log(CP10k + 1) for downstream dimensionality reduction and visualization using Scanpy’s (v1.9.1^57^) *normalize_total* with target sum 10,000 and *log1p*. Gene names and descriptions for Ensembl IDs without annotations were obtained using gget (v0.27.3^58^).

#### Dimensionality reduction and normalization

The concatenated data was mapped to a lower dimensional space by PCA applied to the log-normalized counts filtered for highly variable genes using Scanpy’s *highly_variable_genes*. Next, we computed nearest neighbors and conducted Leiden clustering^59^ using Scanpy.

Initially, this approach was performed on the control and TeNT datasets separately. This resulted in the identification of 19 clusters in the control data and 22 clusters in the TeNT data (Supplementary Figure 11. 16). For both conditions, equal contribution from both datasets indicated that there was minimal batch effect, as expected since the data was sequenced in a pooled sequencing run. We also performed batch correction using scVI^60^ which did not change the contribution of each dataset per cluster. As a result, we continued the analysis using the data that was not batch-corrected with scVI. Next, we concatenated all four datasets and followed the approach described above. This resulted in the identification of 21 Leiden clusters, which we also refer to as cell types (Supplementary Figure 16A). Each cluster was manually annotated with a cell type based on the expression of previously established marker genes^61^. The cell type annotation was validated by the top 20 differentially expressed genes extracted from each cluster using Scanpy’s rank_genes_groups (P values were computed using a t-test and adjusted with the Bonferroni method for multiple testing) (Supplementary Figure 11). Clusters identified as glutamatergic neurons were further broken down into HVC-X- and HVC-RA-projecting glutamatergic neurons using previously established marker genes (data not shown; also see https://github.com/lauraluebbert/TL_2023). We found that reclustering all cells labeled as glutamatergic neurons using the Leiden algorithm did not yield different results and we therefore continued with the initial clusters (data not shown). All results discussed in this paper were confirmed by both jointly and separately clustering the experimental conditions.

#### Comparative analysis of clusters and conditions

Differentially expressed genes between clusters were identified using Scanpy’s *rank_genes_groups* (p values were computed using a t-test and adjusted with the Bonferroni method for multiple testing, and confirmed by comparison to P values generated with Wilcoxon test with Bonferroni correction). In the violin plots, unless otherwise indicated, a star indicates a p value < 0.05 and a fold change > 1.5 difference in mean gene expression between the indicated conditions (p value computed with scipy.stats’ (v1.7.0) *ttest_ind* and adjusted with the Bonferroni method for multiple testing).

### *In situ* hybridization

#### Animals

All of the work described in this study was approved by the California Institute of Technology and Oregon Health & Science University’s Institutional Animal Care and Use Committee and is in accordance with NIH guidelines. Zebra finches (Taeniopygia guttata) were obtained from our own breeding colony or purchased from local breeders. Developmental gene expression in HVC in the 20-, 50-, and 75-days post-hatch (dph) male and female zebra finches was assessed as previously described^62^. The sex of birds was determined by plumage and gonadal inspection. Birds were sacrificed by decapitation, bisected in the sagittal plane and flash-frozen in Tissue-Tek OCT (Sakura-Finetek), and frozen in a dry ice/isopropyl alcohol slurry. Brains of TeNT-manipulated finches were coronally blocked anterior to the tectum and flash frozen in Tissue-Tek (Sakura). All brains were sectioned at 10um on a cryostat and mounted onto charged slides (Superfrost Plus, Fisher).

#### In situ hybridization

*In situ* hybridization was performed as previously described^63,64^. Briefly, DIG-labeled riboprobes were synthesized from cDNA clones for RGS10 (CK312091) and LOC100231469 (class I histocompatibility antigen, F10 alpha chain; DV951963). Slides containing the core of HVC were hybridized overnight at 65°C. Following high stringency washes, sections were blocked for 30 min and then incubated in an alkaline phosphatase conjugated anti-DIG antibody (1:600, Roche). Slides were then washed and developed overnight in BCIP/NBT chromogen (Perkin Elmer). To minimize experimental confounds between animals, sections for each gene were fixed together in 3% paraformaldehyde, hybridized with the same batch of probe, and incubated in chromogen for the same duration.

Sections were imaged under consistent conditions on a Nikon E600 microscope with a Lumina HR camera and imported into ImageJ for analysis. We quantified the expression level of the gene as measured by optical density and the number of cells expressing the gene per unit area, as previously described^62^. Optical density was measured by taking the average pixel intensity of a 300x300 pixel square placed over the center of HVC. This value was normalized to the average background level of the tissue. To quantify the number of labeled cells, we established a threshold of expression that was 2.5x the background level. Binary filters (Close-, Open) were applied and the number of particles in the same 300x300 pixel square was quantified.

### Histology

After cardiac perfusion with room temperature 3.2% PFA in 1xPBS we let the brains fix for 2-4 hours at room temperature. After each hemisphere of the brain was sectioned sagittally with a vibratome at 70-100 µm thickness. The brain slices containing HVC were collected and incubated at 4 C overnight with the primary rabbit anti-GFP (AB3080P, EMD Milipore) (blocked in 10% donkey serum in 0.2% Triton 1xPBS). On the second day, the brains were washed in 0.05% Triton 1xPBS and incubated for 2 hours in the dark at room temperature in the secondary goat anti-rabbit 488 (ab150077). Next, the brain slices were washed and mounted in Fluoromount (Sigma). Confocal images were taken with the LSM800.

## Data and Code Availability

Data generated in this study have been deposited in Caltech DATA and can be found at the following DOIs: https://doi.org/10.22002/ednra-nn006 and https://doi.org/10.22002/3ta8v-gj982. Please do not hesitate to contact the authors for data or code requests. The code used for the analysis of the single-cell RNA sequencing data can be found here: https://github.com/lauraluebbert/TL_2023. The code used for the analysis of the chronic electrophysiology data can be found here: https://github.com/jordan-feldman/Torok2023-ephys.

## Author contributions

Conceived the ideas of experimental design of the study – ZT, CL

Perform behavioral, single-cell, and electrophysiology experiments/data collection – ZT

Perform molecular or imaging experiments/data collection – ZT, LL, SW, AAN, JP (from methods)

Data analysis and interpretation – ZT, LL, JF, AD, WG

Manuscript writing – ZT, CL, AD, AF, LL, LP

Stylistic/grammatical revisions – All authors

Provided revisions to the scientific content of manuscript - All authors

Provided funding – ZT, AF, LP, CL

## Acknowledgments

Funding source: R01NS104925A (CL, AF), Chen Graduate Innovator Grant 2019 (ZT) We thank Jeff Park and the Caltech Sequencing Facility for the library generation.

We thank Daniel Pollak for early chronic electrophysiology advice and Allan Hermann-Pool for advice on the single-cell RNA sequencing experimental pipeline.

## Supplementary Figures

**Supplementary Figure 1:**
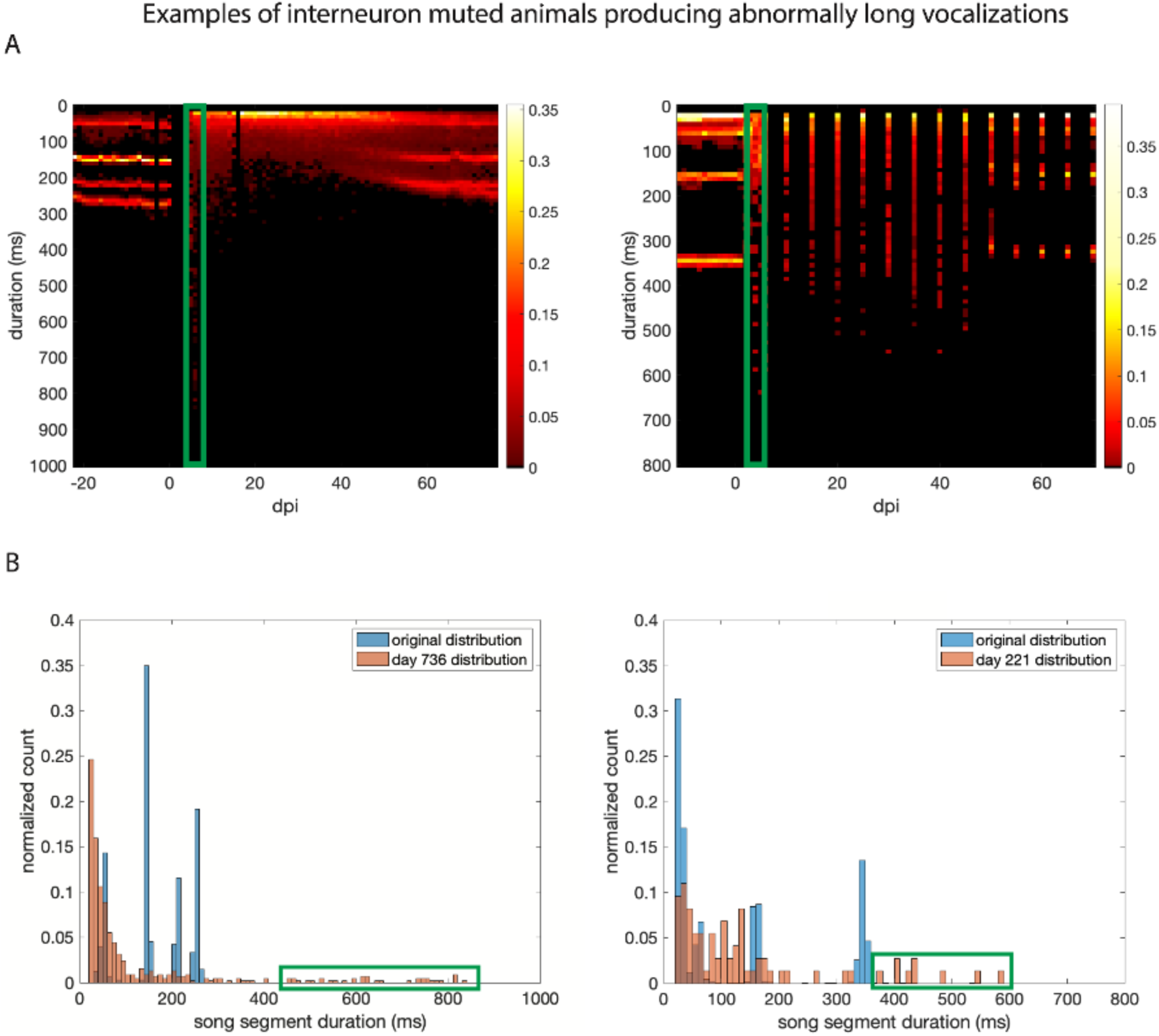
Examples of abnormally long syllable lengths after injection of interneuron muting virus in two animals. **a** Syllable length durations for the length of song degradation and recovery. The Y axis depicts the length of the syllables in milliseconds plotted over days post-injection (dpi) of either TeNT virus. TeNT-treated animals displayed a short period during which some vocalizations were of length not observed in normal animals and eventually became highly variable and shorter (shifts to shorter length sounds). The green rectangle highlights the day post-injection portrayed in B for each animal. **B** Histogram of syllable durations (blue trace is before injection of virus, orange trace is after injection of virus). The green rectangle highlights vocalizations of abnormal length.

**Supplementary Figure 2:**
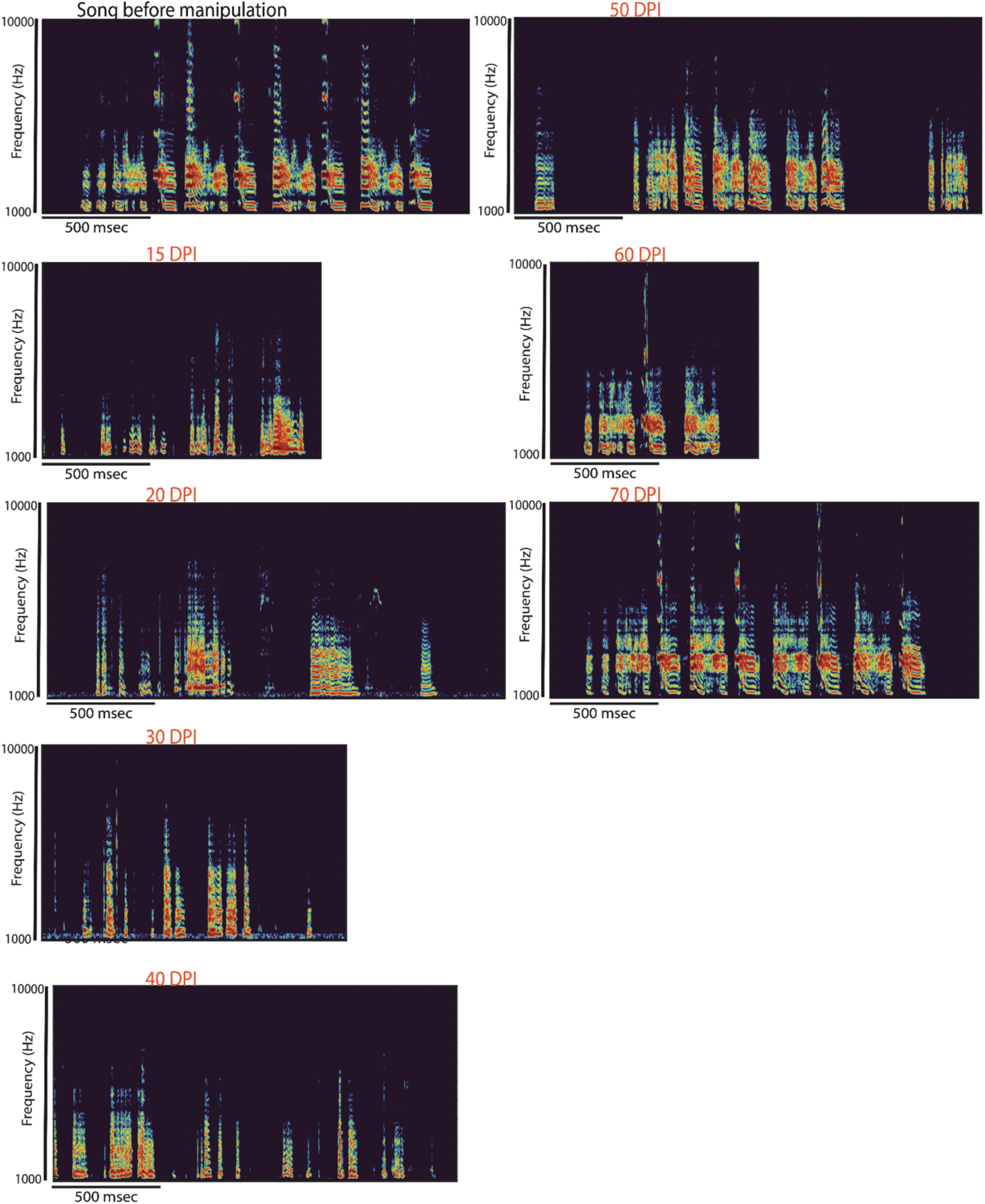
Example spectrograms of a TeNT-treated animal (B138) during song degradation and recovery. Vocalizations between 15 dpi and 30 dpi were much shorter than the first long syllables shown in Figure 1 A at 5 dpi.

**Supplementary Figure 3:**
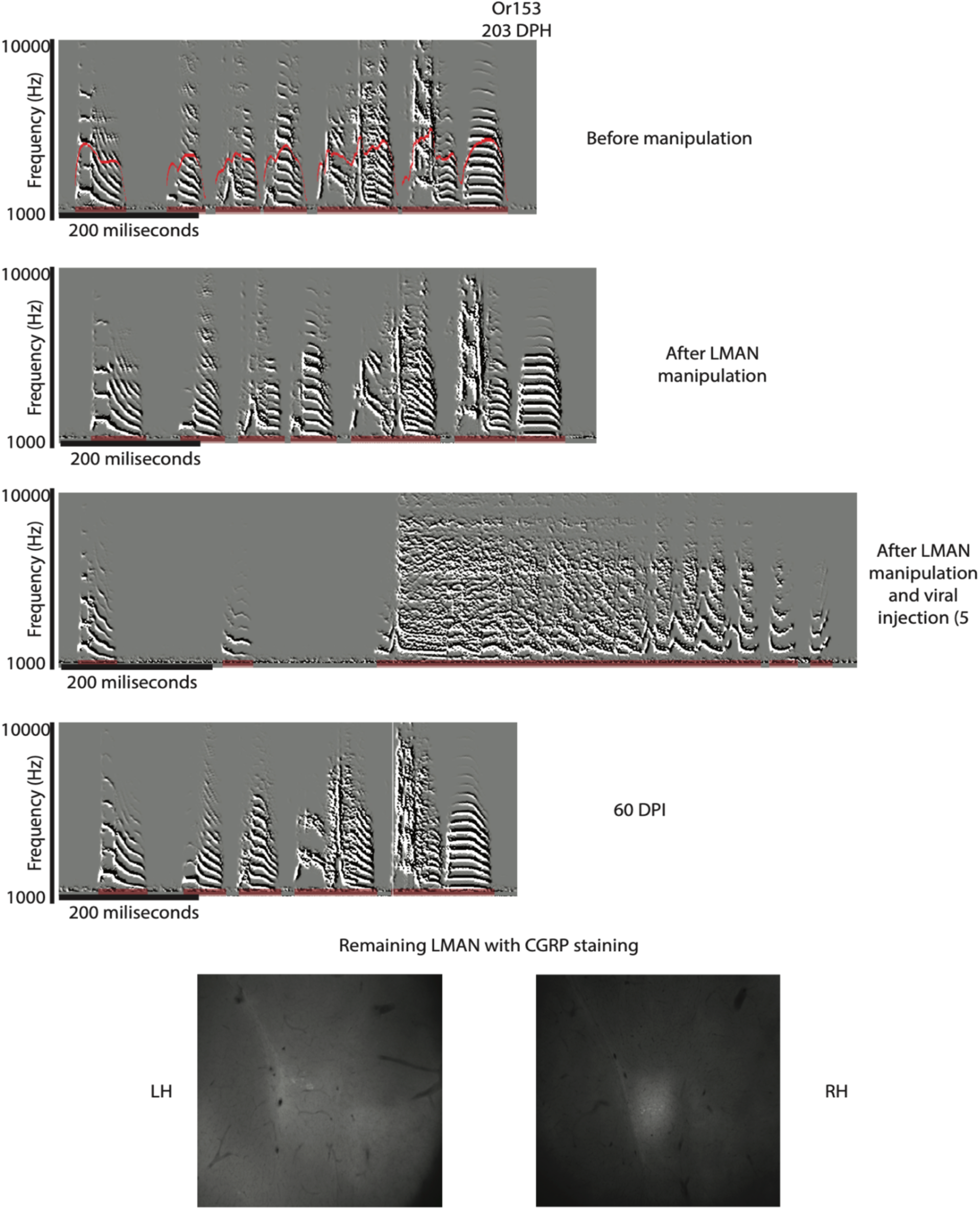
Song degradation and recovery after chronic removal of inhibition in an animal without LMAN. Spectrograms are showing the song of the animal before and after LMAN lesion at 5 and 40 days post viral injection (dpi). The histology image shows the amount of LMAN left (based on CGRP staining) in the right hemisphere (RH).

**Supplementary Figure 4:**
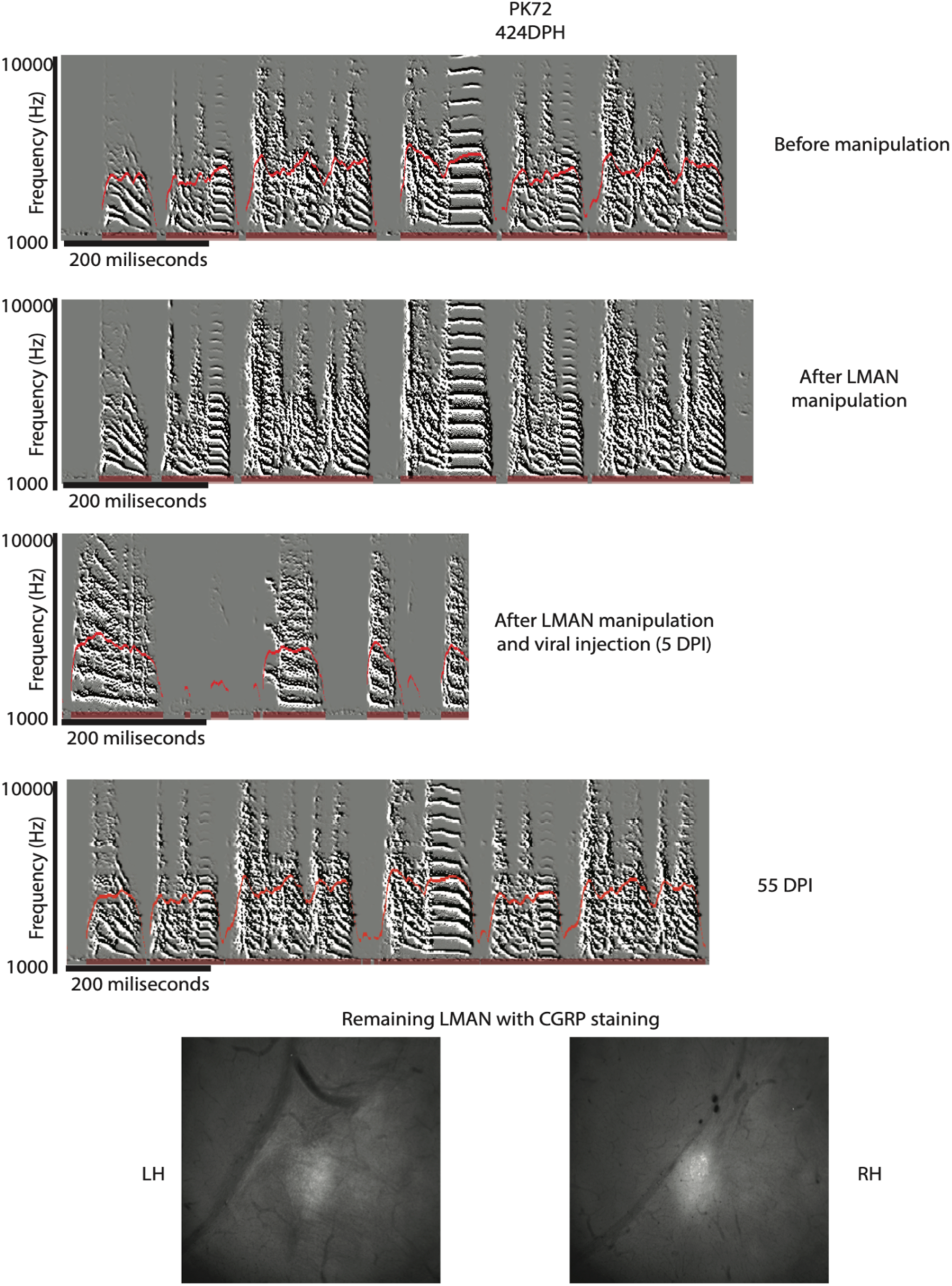
Song degradation and recovery after chronic removal of inhibition in an animal without LMAN. Spectrograms are showing the song of the animal before and after LMAN lesion at 5 and 60 days post viral injection (dpi). The histology images indicate the amount of LMAN left (based on CGRP staining) in the left (LH) and right (RH) hemispheres.

**Supplementary Figure 5:**
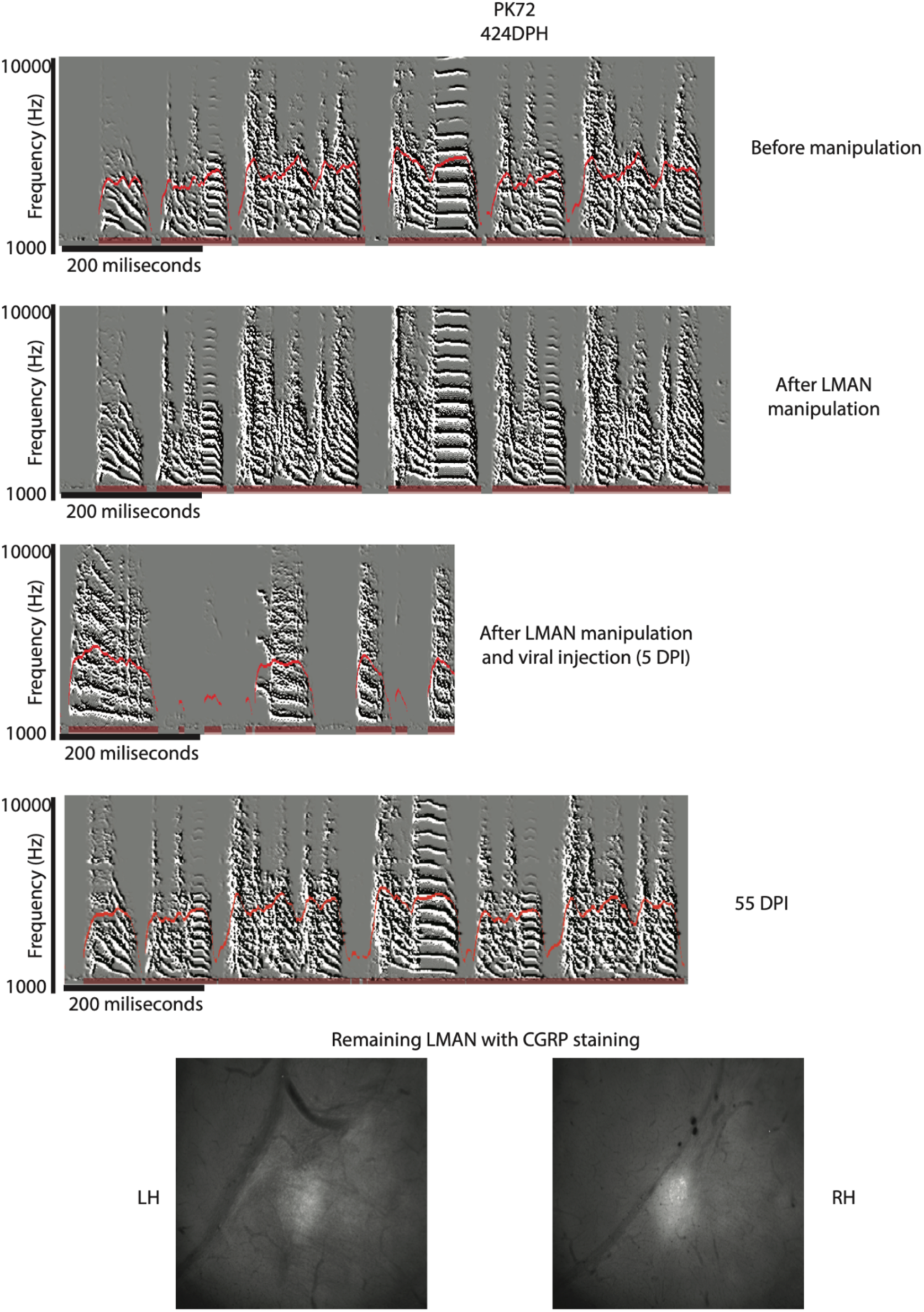
Song degradation and recovery after chronic removal of inhibition in an animal without LMAN. Spectrograms are showing the song of the animal before and after LMAN lesion at 5 and 55 days after viral injection (dpi). The histology images indicate the amount of LMAN left (based on CGRP staining) in the left (LH) and right (RH) hemispheres.

**Supplementary Figure 6:**
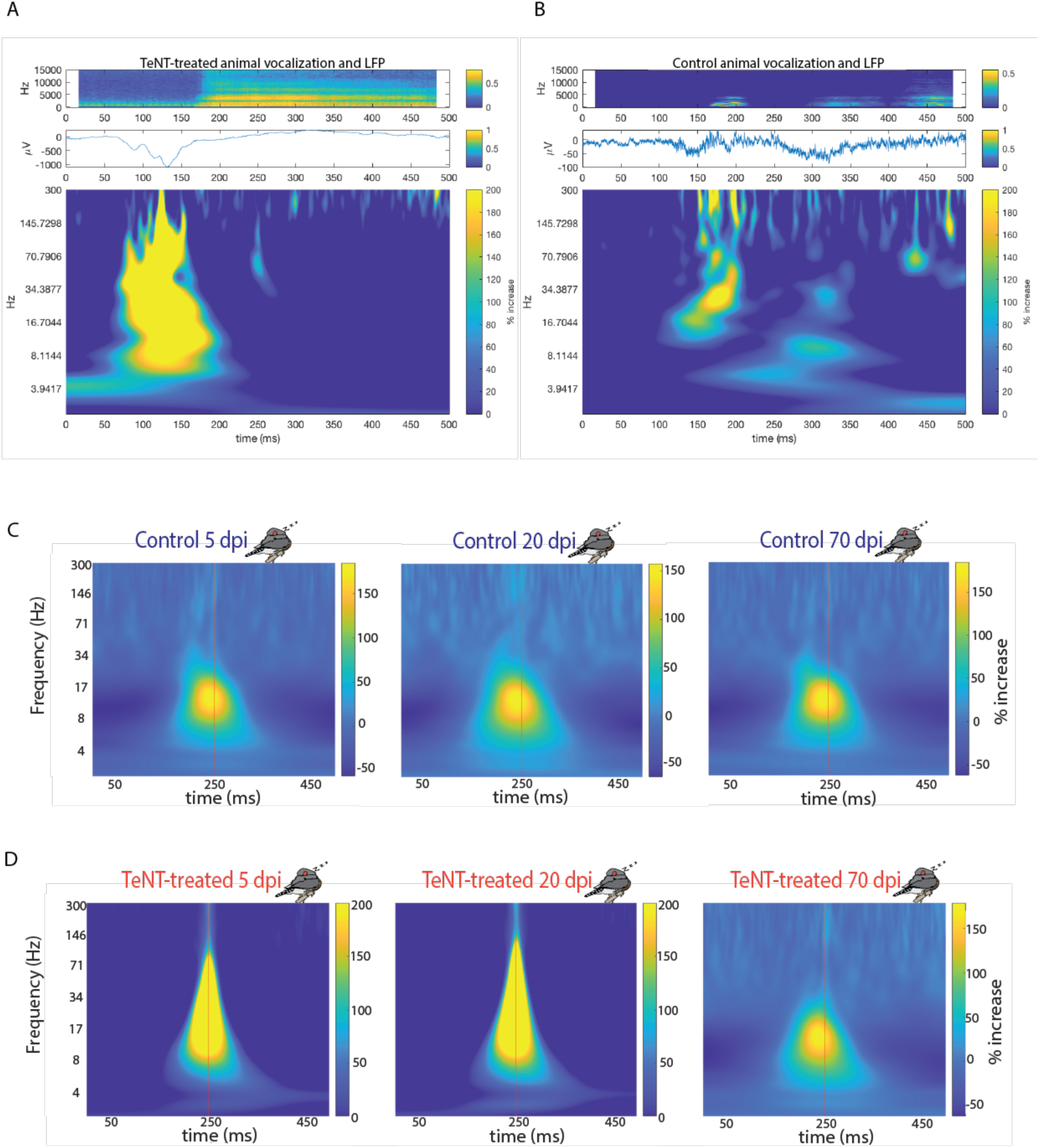
Example electrophysiology traces averaged over 5 instances of normal or 5 instances of degraded vocalizations and averages of sleep voltage deflections. **a** Averaged spectrogram of degraded vocalization (n=5) 5 days post-electrode-implantation in a chronically recorded TeNT-treated animal. The plot below the spectrogram shows the raw averaged trace of extracellular recording. Below the raw trace is the averaged continuous wavelet transform of the local field potentials (LFP, 1-300Hz). The plots show a large deflection event (similar to those seen during lights-off in Figures 2 and 3) right before the onset of the vocalization in the TeNT-treated animal. **B** Averaged song spectrogram (n=5) 5 days post-electrode-implantation from a chronically recorded control animal. The plot below the spectrogram shows the raw averaged trace of extracellular recordings. Below the raw trace is the averaged continuous wavelet transform of the local field potentials (LFP, 1-300 Hz). The averaged control song shows more and smaller amplitude deflections mostly during the vocalization compared to the TeNT-treated vocalization. **C-D** Spectral decomposition of local field potentials (LFPs) of the averaged deflections during night time at 5, 20, and 60 dpi in one control (C) and one TeNT-treated animal (D). The vertical red line depicts the trough of the raw deflection trace. The % increase is the relative increase compared to non-deflection events in the same recording timeframe of the same animal (Methods). Deflections of TeNT-treated animals in the 15-30 Hz range were approx. 27 times larger than those in control animals at 5 dpi. These differences are statistically significant between control and TeNT-treated groups, but not within the control group (p=0.7631 between controls, p<10^-35^ between all other pairs). The deflections across all animals and frequencies become more similar by 60 dpi (15-30 Hz: p=0.1371 between controls, p<10^-6^ between all other pairs; 30-70 Hz: p=0.7493 between controls, p<10^-2^ between all other pairs). For details on statistics see Methods.

**Supplementary Figure 7:**
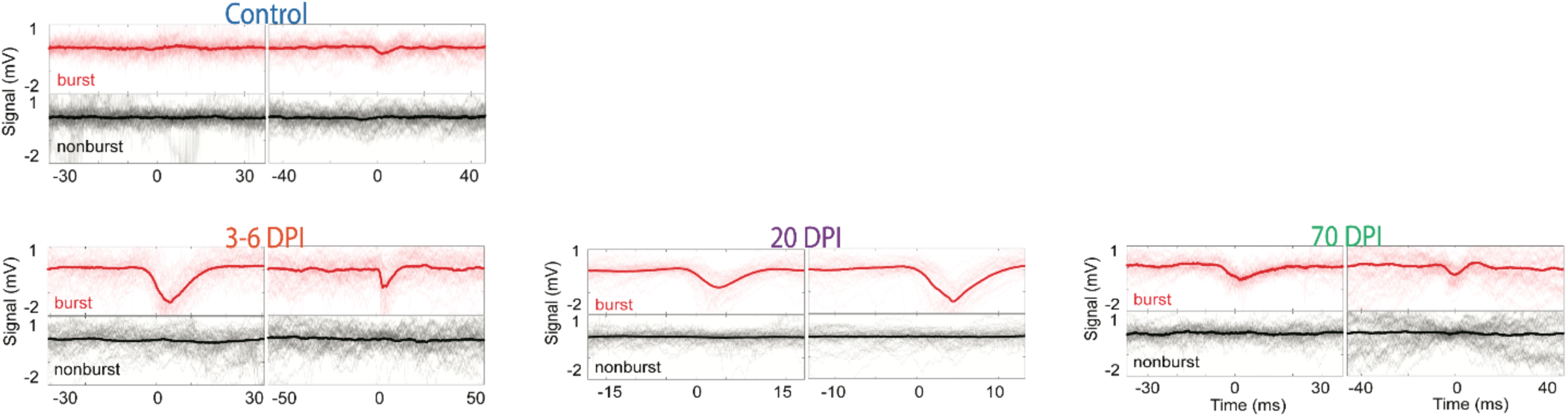
Example traces of raw deflections in the acute Neuropixel recordings during lights-off periods. Control animals barely showed any visible deflection events, while TeNT-treated animals (example traces shown at 3-6, 20, and 70 dpi) displayed large amplitude voltage deflection events.

**Supplementary Figure 8:**
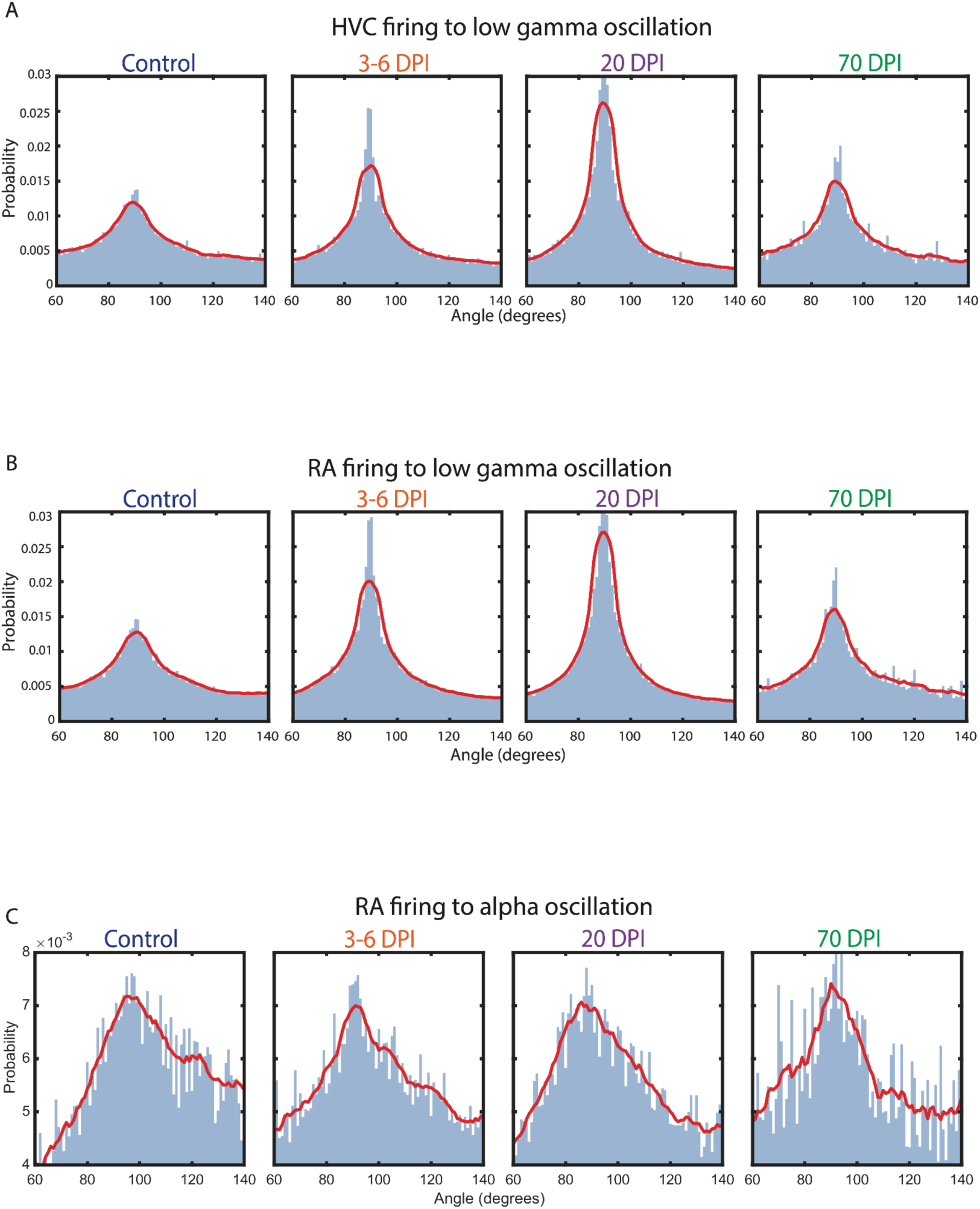
Relationship between alpha or gamma oscillations during lights-off voltage deflection events and local neuronal firing in HVC and RA from the acute NPIX recordings. **a** The normalized probability distribution of neurons locally within HVC fire during a specific phase (angle) of the gamma (30-40 Hz) oscillations extracted from the LFP signal of the averaged deflection events at 3-6 dpi (n=4 animals), 20 dpi (n=4 animals), 70 dpi (n =2 animals that recovered their song by then). There was a slight change in local neuronal firing to the angle and locking precision to gamma oscillations that resembled control by 70 dpi. **B** Normalized probability distribution of neurons locally within RA fire during a specific phase (angle) of the alpha (1-10 Hz) oscillations in HVC extracted from the LFP signal of the averaged deflection events at 3-6 dpi, 20, 70 dpi. No change in RA spontaneous neuronal firing to alpha oscillations in HVC during the deflection events over the course of the manipulation. **C** Normalized probability distribution of neurons locally within RA fire during a specific phase (angle) of the gamma (30-40 Hz) oscillations in HVC extracted from the LFP signal of the averaged deflection events at 3-6, 20, and 70 dpi. We observed no change in RA spontaneous neuronal firing to the gamma oscillations in HVC during the deflection events over the course of the manipulation.

**Supplementary Figure 9:**
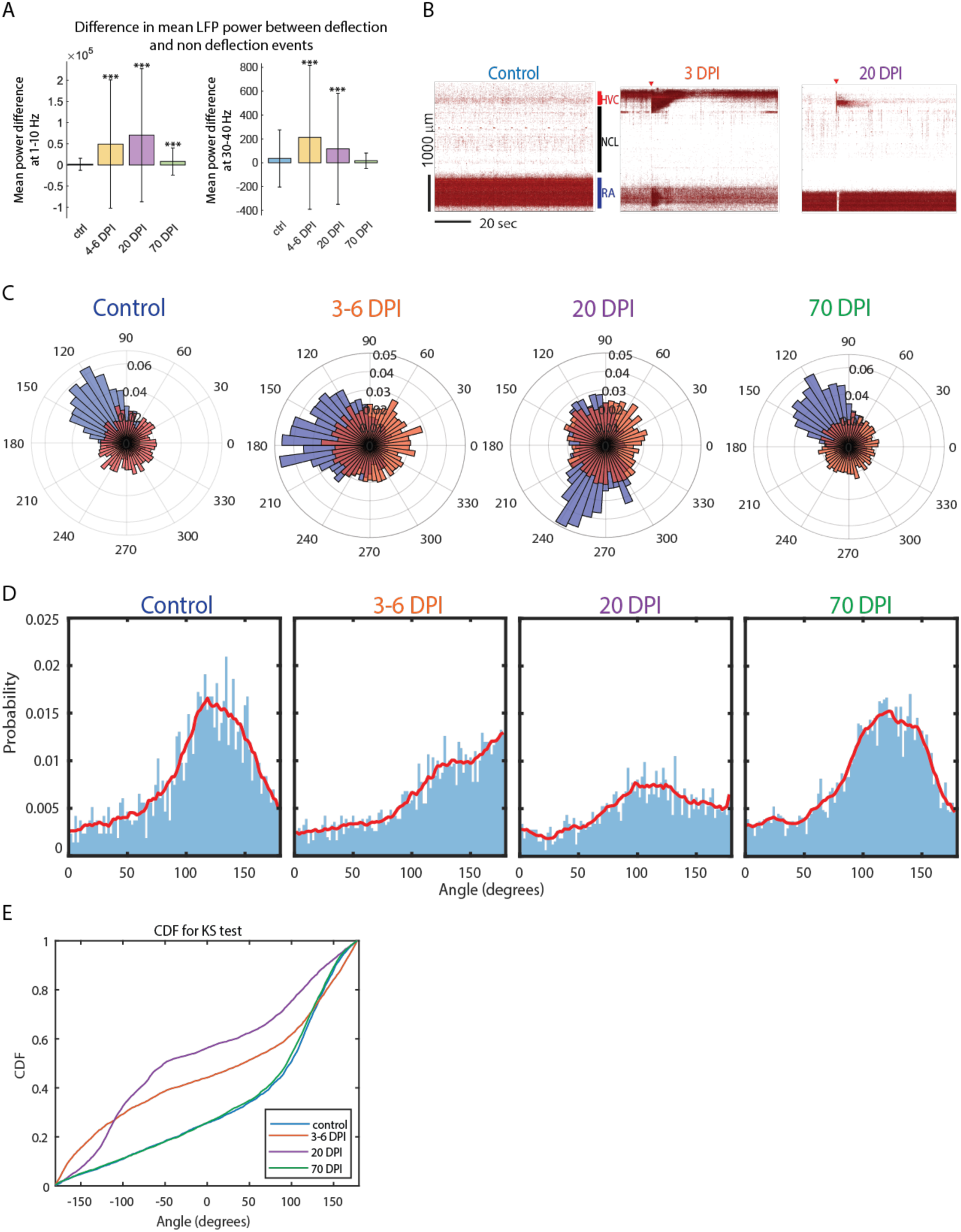
Quantification of the angle relationship between alpha (1-10 Hz) and gamma (30-40 Hz) frequencies during deflection events in control and TeNT-treated animals during acute head-fixed recordings. **A**: The average difference in power (at alpha, 1-10 Hz, and low gamma 30-40 Hz frequency ranges) between voltage deflection and non-deflection events in control, 3-6, 20, 70 dpi. The power content in the alpha range increased in a statistically significant way (Wilcoxon, rank sum test) between control and 3-6 (p=2.9*10^-36), 20 (p=8.6*10^-167), 70 (p=5.4*10^-4) dpi animals. However, the increase in power between control and 3-6 (p=2.2*10^-37), 20 (p=6.3*10^-27) dpi is statistically significant but returns to control level by 70 (p=0.37) dpi. The stars above the bar plots (*) indicate statistical significance (*: p < 0.005, **: p< 0.01, ***: p < 0.001). **B** Examples of neuronal activity in a control animal, TeNT-treated animals at 3 and 20 dpi. The red arrows highlight the “superbursts” or extreme firing levels within HVC and RA which we observed in 3 animals (two animals at 3-4 dpi and one at 20 dpi) in a total of seven instances.**C**: The polar histograms of the angle of the alpha oscillations (1-10Hz) at the maximum amplitude of the gamma oscillation (30-40 Hz) during deflection events. The red distribution represents a randomly shuffled dataset, while the blue is the true distribution of angles in control (n=3), and TeNT-treated animals at 3-6 dpi (n=4), 20 dpi (n=4) and 70 dpi (n=4) during deflection events. **D**: Relationship of alpha and low gamma oscillations during deflection events in control (n=3), 3-6 (n=4), 20 (n=4), and 70 (n=2 animals) dpi animals (over animals and conditions). The probability distribution of a specific angle of the low-frequency oscillation at the maximum amplitude of the gamma oscillation. **E**: The results of the Kolgomorov-Smirnov test on the cumulative density function (CDF) to assess if the change in probability distribution shown in C is statistically significant from control distributions at 3-6, 20, and 70 dpi. The purple (20 dpi) and orange (3-7 dpi) distributions differ significantly from the blue control and the 70 dpi green distributions. The 70 dpi population is not significantly different from the control group.

**Supplementary Figure 10:**
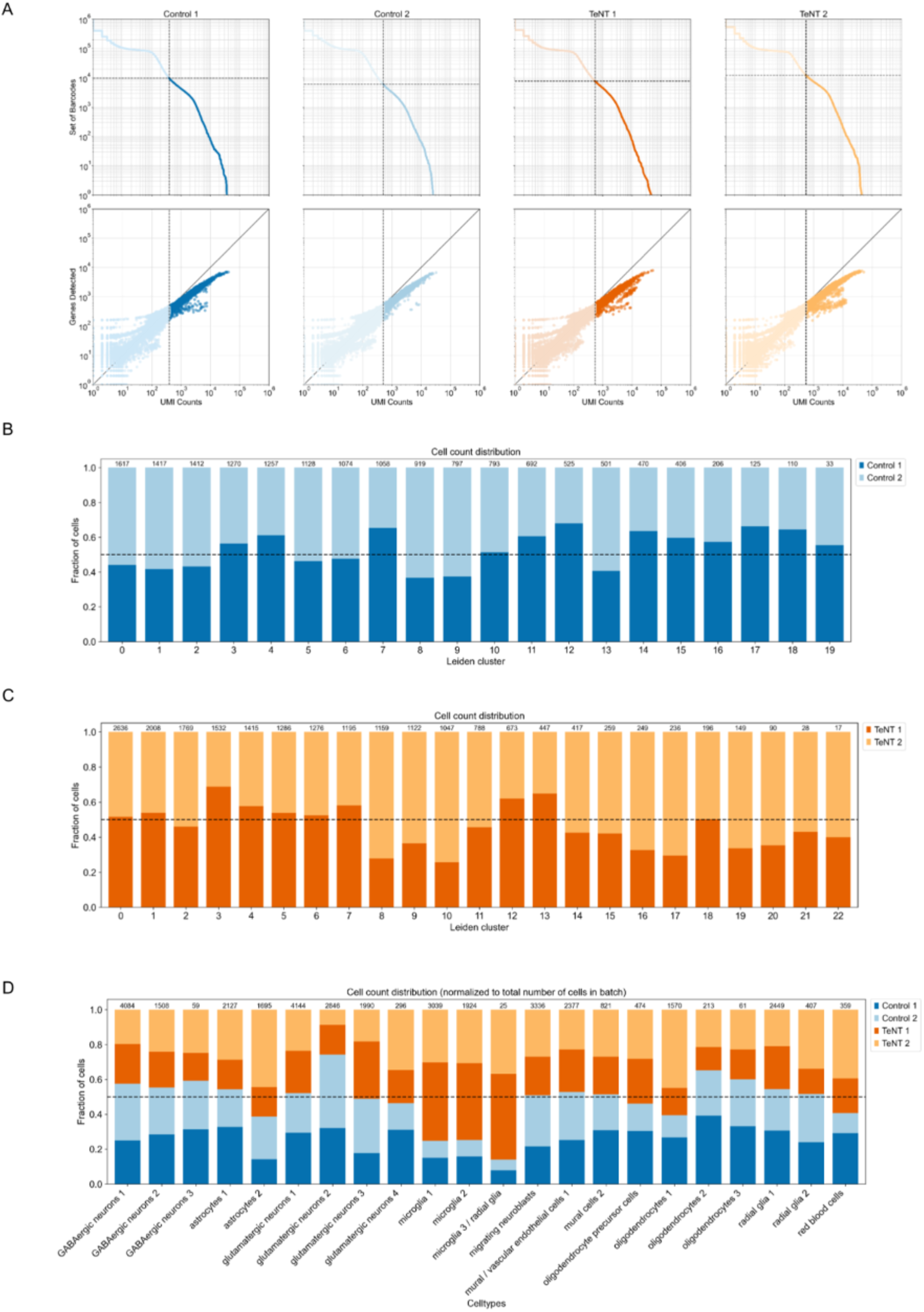
Quality control of the single-cell RNA sequencing HVC datasets from control and TeTN-treated animals at 25 days post-injection (dpi). **a** “Knee plots” showing the set of barcodes (top row) and number of genes detected (bottom row) over UMI counts. The dashed lines depict the quality filtering cutoff. **B-C** Barplot depicting the fraction of cells from each replicate per cluster for control (B) and TeNT (C), normalized (by dividing) to the total number of cells in each replicate. Control and TeNT datasets were clustered separately using the Leiden algorithm. The equal distribution of replicates across the clusters suggests that technical effects do not dominate the clusters. Thus, we did not perform batch correction. The numbers on top of the bars indicate the total number of cells in each cluster. **D** Barplot depicting the fraction of cells from each dataset in the cell type clusters obtained after jointly clustering the control and TeNT datasets. The numbers on top of the bars indicate the total number of cells in each cluster.

**Supplementary Figure 11:**
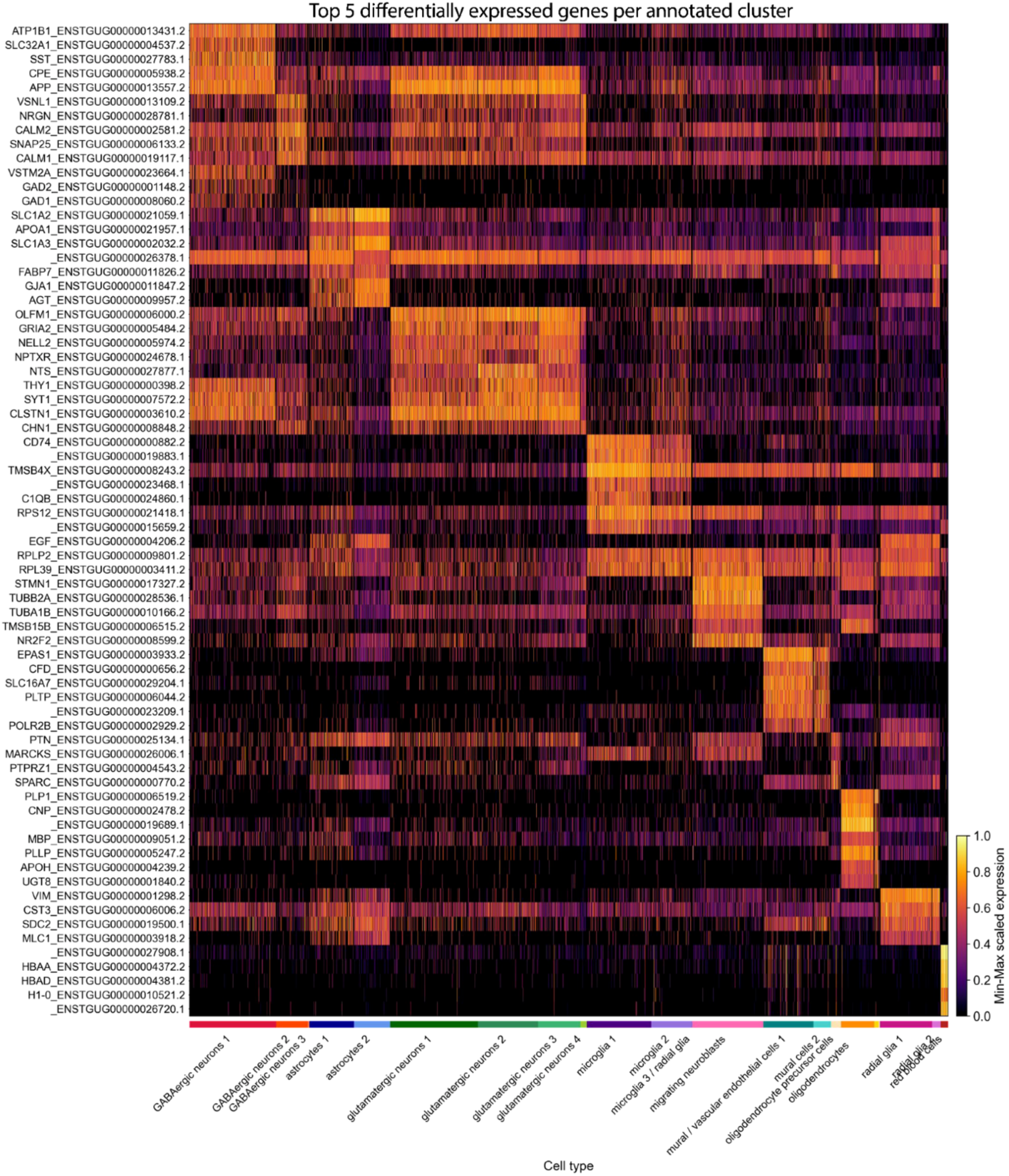
Heatmap of top 5 differentially expressed genes per annotated cell type/cluster obtained by single-cell RNA sequencing of HVC from control and TeNT-treated birds at 25 dpi. Differentially expressed genes between clusters were identified using Scanpy’s rank_genes_groups (p values were computed using a t-test and were adjusted with the Bonferroni method for multiple testing. They were then confirmed by comparison to p values generated with the nonparametric Wilcoxon test with Bonferroni correction). The heatmap depicts the min-max scaled expression for each gene.

**Supplementary Figure 12:**
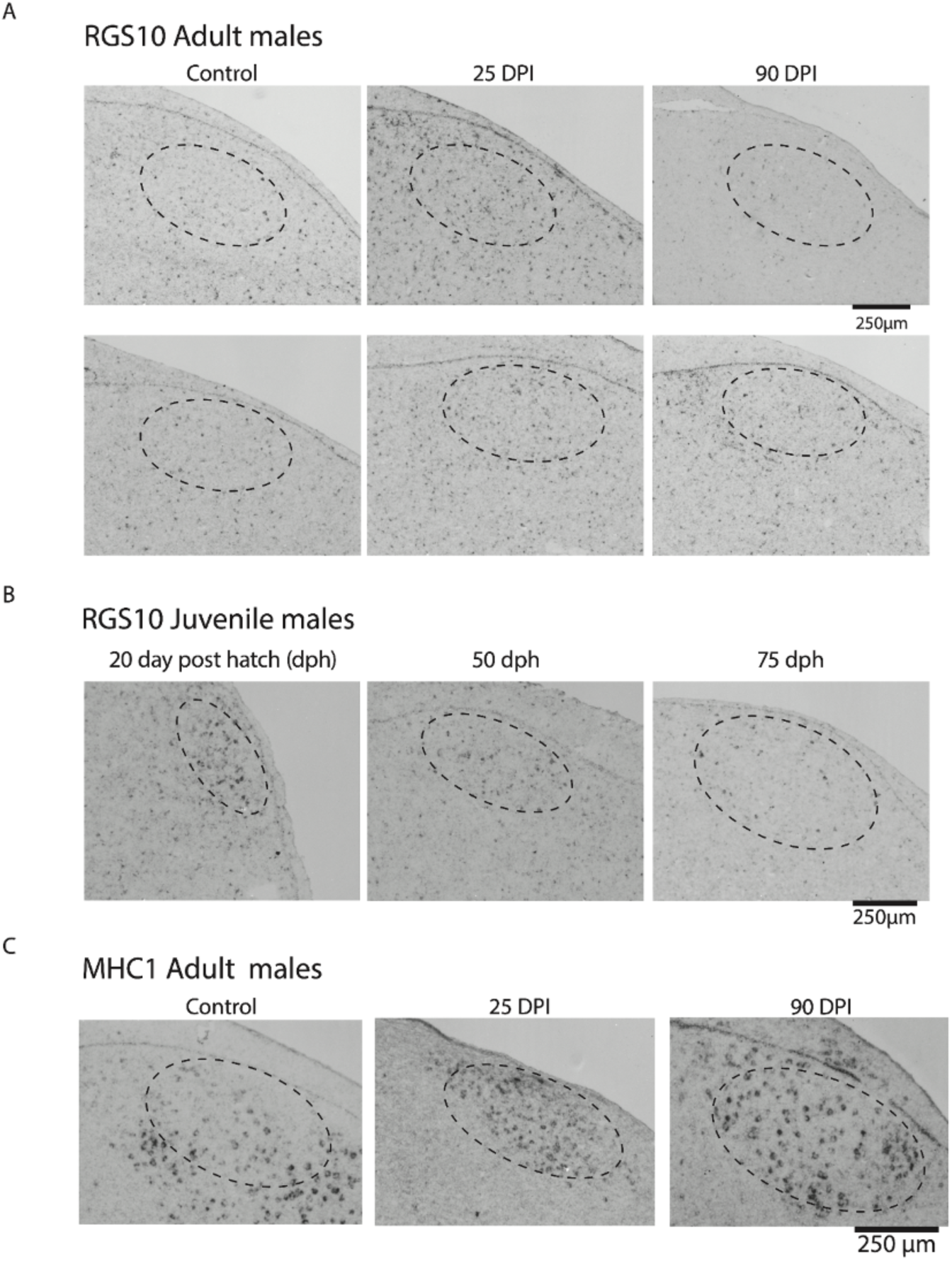
In situ hybridization of microglia marker gene RGS10 in adult male control, TeNT-treated and juvenile male HVC & MHC1 gene in adult male control and TeNT-treated HVC. **a** Histological sections of HVC (in control and TeNT-treated animals at 25 and 90 dpi) after in situ hybridization of RNA probes for RGS10 (a gene marker for microglia). **B** Histological sections of HVC in naive juvenile males (at 20, 50, and 75 days post-hatching (dph)) after in situ hybridization of RNA probes for RGS10. **C** Histological sections of HVC (from control and TeNT-treated animals at 25 and 90 dpi) after in situ hybridization of RNA probes for MHC1. Black/darker dots indicate enzyme reactions resulting in successful probe localization and suggest target gene expression.

**Supplementary Figure 13:**
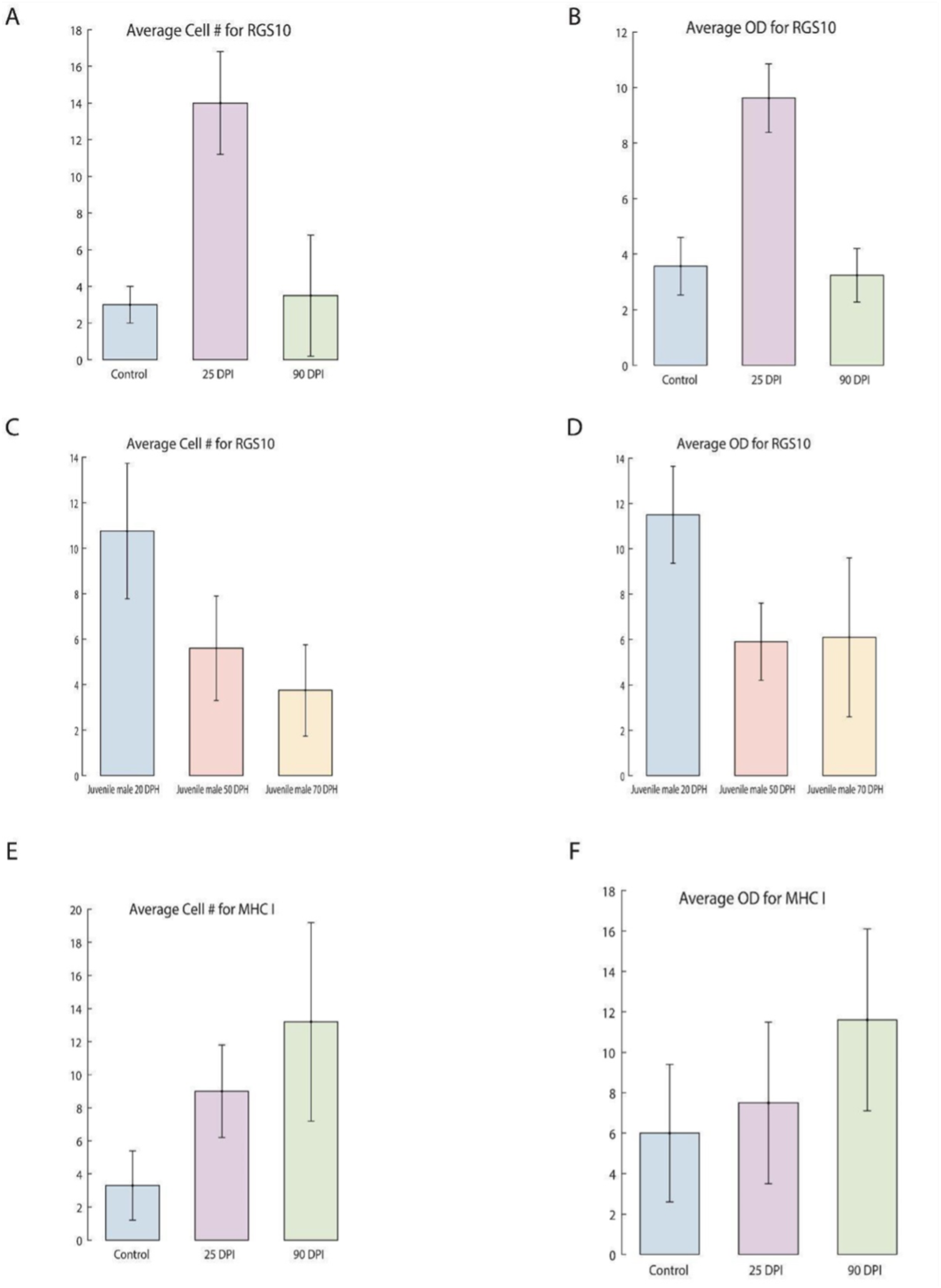
Quantification of the in situ hybridization against microglia marker gene RGS10 in adult male control, TeNT-treated and juvenile male HVC; and MHC1 in adult male control, TeNT-treated animals. **A-B** Quantification of the in situ hybridization for RGS10 between control (n=4 animals) and TeNT-treated animals at 25 dpi (n=4) and 90 dpi (n=4). **C-D** Quantification of the in situ hybridization for RGS10 between juvenile males at 20, 50, and 70 days post-hatching (dph) (n=4). **E-F** Quantification of the in situ hybridization for MHC1 between control (n=4) and TeNT-treated animals at 25 (n=4) and 90 dpi (n=4). Error bars represent standard deviation.

**Supplementary Figure 14:**
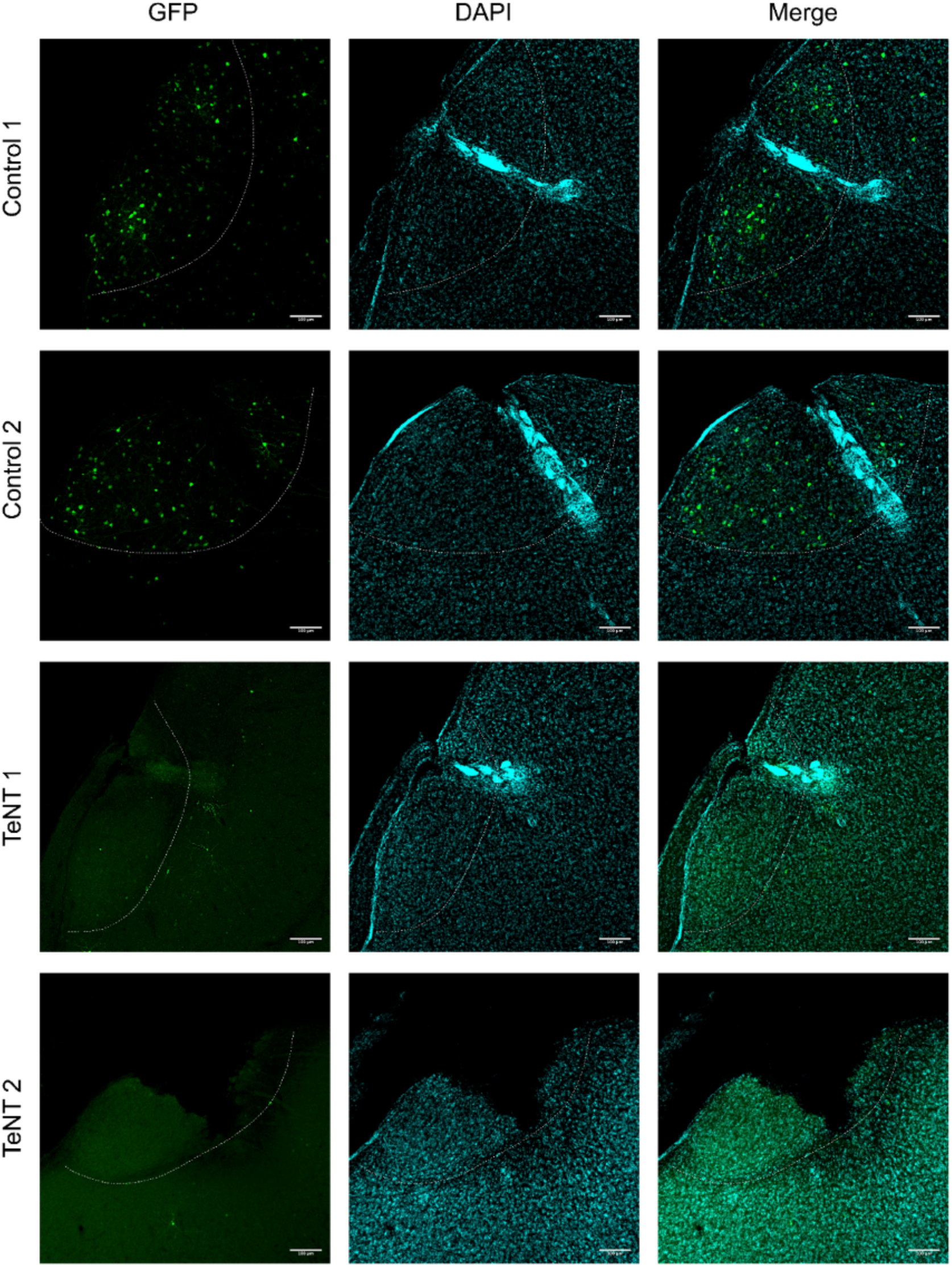
Histology of electrode array location in HVC in the chronically implanted animals. The white dotted line outlines HVC. Some sections display missing tissue due to the removal of the electrodes after perfusion of the animals. The stronger cyan signal indicates glial scar formation around the electrode array, which provides an approximation of the location of the electrodes. Electrodes located closer to the bottom of HVC close to the shelf were not used for analysis.

**Supplementary Figure 15:**
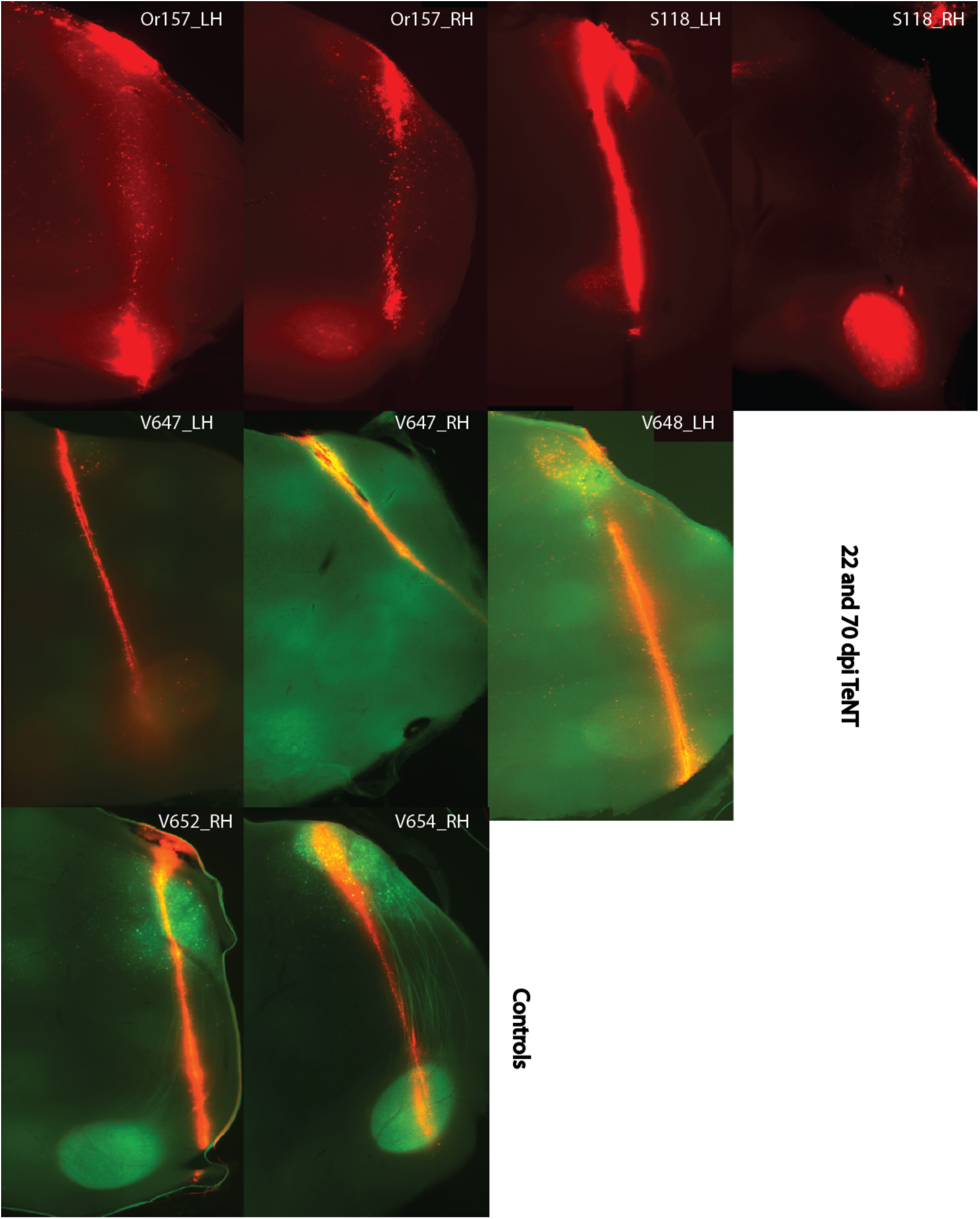
Histology to confirm the high-density silicone electrode location in the acute head-fixed animal recordings. The red trace represents the electrode location. The green trace represents the second electrode location in animals that were recorded twice, 40 days apart. The white labels represent the animal IDs. “LH” and “RH” stands for left and right hemisphere, respectively.

**Supplementary Figure 16:**
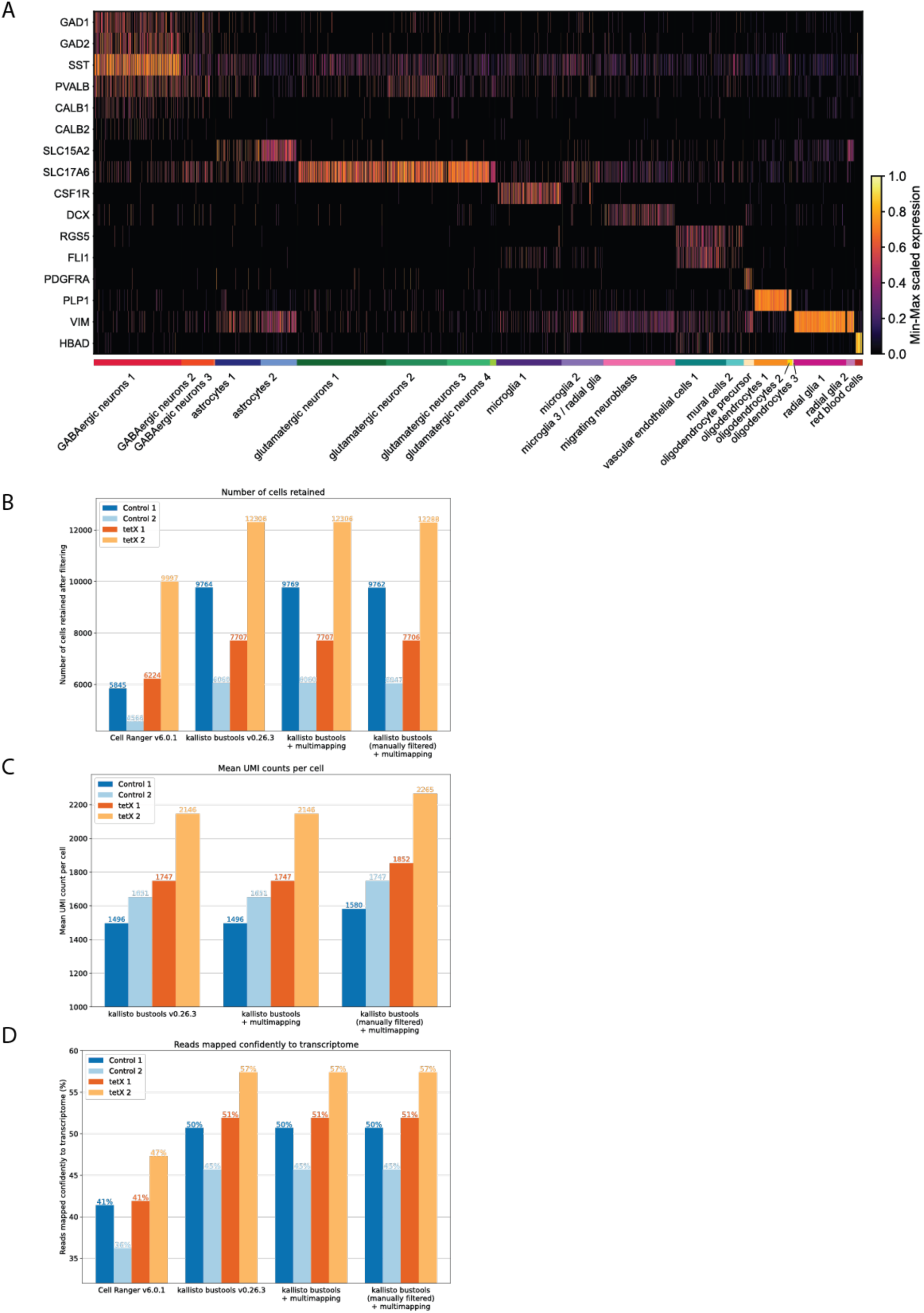
Comparison of different pre-processing methods for the HVC single-cell RNA sequencing datasets and marker genes. **a** Heatmap showing min-max scaled expression of cell type marker genes for each cell type (data from both control and TeNT-treated animals). **B** Number of cells retained after quality control for each dataset and alignment method. **C** Mean UMI counts per cell for each dataset and pre-processing method. **D** Percentage of reads confidently mapped to transcriptome for each pre-processing method.

**Supplementary Table 1:** Amplitudes and durations of the chronic voltage deflections measured throughout the recording. **A** Mean duration (calculated as the distance from the onset to the half-width point of the event) in ms of voltage deflection events with standard deviation, each row represents an event from one control (Or 295, PK31) or TeNT-treated (B138, Or296) animal. The data was sampled at 3-5, 15, 30, 45, 60, and 75 dpi. **B** Mean amplitudes (in µV) of voltage deflection events with standard deviation.

| Deflection durations (half-widths) in ms (mean ± SD) | | | | | | |
| --- | --- | --- | --- | --- | --- | --- |
|  | 3-5 DPI | 15 DPI | 30 DPI | 45 DPI | 60 DPI | 75 DPI |
| Control animal 1 (OR295) | 44.2083<br>$\pm 16.0885$ | 50.7321<br>$\pm 21.5322$ | 47.7079<br>$\pm 16.4028$ | 49.5889<br>$\pm 18.5678$ | 45.4170<br>$\pm 15.4166$ | 48.2883<br>$\pm 18.5856$ |
| Control animal 2 (PK31) | 43.9120<br>$\pm 15.5607$ | 46.5072<br>$\pm 17.6237$ | 36.7974<br>$\pm 15.6453$ | 42.4989<br>$\pm 15.8985$ | 37.8048<br>$\pm 15.2497$ | 35.3157<br>$\pm 15.2297$ |
| TeNT-treated animal 1 (B138) | 24.7074<br>$\pm 11.0296$ | 20.4568<br>$\pm 5.7482$ | 26.2938<br>$\pm 10.4427$ | 32.1460<br>$\pm 13.1017$ | 36.8212<br>$\pm 15.3777$ | 41.4122<br>$\pm 15.9937$ |
| TeNT-treated animal 2 (OR296) | 34.6744<br>$\pm 10.7396$ | 29.2852<br>$\pm 8.3464$ | 29.5713<br>$\pm 9.5446$ | 31.4171<br>$\pm 11.5461$ | 39.8151<br>$\pm 10.5067$ | 42.9507<br>$\pm 14.0682$ |

**Supplementary Table 1: Amplitudes and durations of the chronic voltage deflections measured throughout the recording.**
| Deflection amplitudes in μV (mean ± SD) | | | | | | |
| --- | --- | --- | --- | --- | --- | --- |
|  | 3-5 DPI | 15 DPI | 30 DPI | 45 DPI | 60 DPI | 75 DPI |
| Control animal 1 (OR295) | -136.3911<br>$\pm 28.7491$ | -155.0317<br>$\pm 49.3434$ | -151.5708<br>$\pm 35.8116$ | -149.0358<br>$\pm 34.9365$ | -152.4109<br>$\pm 33.1907$ | -150.5303<br>$\pm 33.8846$ |
| Control animal 2 (PK31) | -111.9867<br>$\pm 20.8258$ | -117.1694<br>$\pm 26.5227$ | -133.9981<br>$\pm 26.5174$ | -130.6166<br>$\pm 25.9310$ | -138.6591<br>$\pm 26.7510$ | -135.2756<br>$\pm 25.9969$ |
| TeNT-treated animal 1 (B138) | -538.6617<br>$\pm 307.8410$ | -816.7580<br>$\pm 371.9481$ | -705.8811<br>$\pm 267.1738$ | -422.1097<br>$\pm 143.1100$ | -306.8821<br>$\pm 101.9335$ | -261.2531<br>$\pm 80.5379$ |
| TeNT-treated animal 2 (OR296) | -174.6613<br>$\pm 39.6156$ | -251.7470<br>$\pm 70.1546$ | -337.3158<br>$\pm 94.5369$ | -525.5973<br>$\pm 157.8933$ | -228.3434<br>$\pm 48.9163$ | -162.6949<br>$\pm 35.6879$ |

**Supplementary Table 2:** Overview of single-cell RNA sequencing datasets.

| Dataset (short name) | Species | Condition | Brain area | Replicate # | Technology | Pre-processing tool | # of cells retained after QC | Sequencing depth (number of reads processed) | Reads mapped confidently to transcriptome (%) | Mean UMI count per cell | Total UMI count |
| --- | --- | --- | --- | --- | --- | --- | --- | --- | --- | --- | --- |
| C1 | Taeniopygia guttata | cag-neonGreen | HVC (both hemispheres) | 1 | 10xv3 | kallisto bustools | 9,763 | 744,473,151 | 50.7 | 1580.8531 | 15,433,015 |
| C2 | Taeniopygia guttata | cag-neonGreen | HVC (both hemispheres) | 2 | 10xv3 | kallisto bustools | 6,047 | 787,232,472 | 45.7 | 1747.639 | 10,568,132 |
| E1 | Taeniopygia guttata | dlx-TeNT-GFP | HVC (both hemispheres) | 1 | 10xv3 | kallisto bustools | 7,706 | 867,768,600 | 51.9 | 1852.4215 | 14,274,784 |
| E2 | Taeniopygia guttata | dlx-TeNT-GFP | HVC (both hemispheres) | 2 | 10xv3 | kallisto bustools | 12,288 | 810,253,355 | 57.4 | 2265.4258 | 27,837,586 |

